# Olfactory Deficits and Structural Anomalies of the Peripheral Olfactory System in 22q11.2 Deletion Syndrome

**DOI:** 10.64898/2026.10.05.756775

**Authors:** David R. Roalf, Donna M. McDonald-McGinn, Ally Atkins, Monica E. Calkins, T. Blaine Crowley, Phoebe Freedman, R. Sean Gallagher, Christina Mastracchio, Daniel E. McGinn, Austin Moran, Audrey Olson, Maggie K. Pecsok, Lydia Rockart, Oanh Tran, Elaine H. Zackai, Beverly S. Emanuel, Kosha Ruparel, Russell T. Shinohara, Lauren K. White, J. Eric Schmitt, Raquel E. Gur

## Abstract

**Background:** Approximately 25% of individuals with 22q11.2 deletion syndrome (22q11DS) develop a psychotic disorder, and the syndrome shares key neurodevelopmental features with psychosis in the general population. The olfactory system develops early in utero, providing a window into gestational disruptions relevant to psychosis risk. We hypothesized that 22q11DS would show impairments in olfactory function and structural anomalies of the olfactory system relative to psychosis spectrum (PS) youth without a known genetic syndrome and typically developing controls (TD).

**Methods:** Participants included individuals with 22q11DS (n=101), PS (n=56), and TD (n=52). Sinonasal symptoms, odor identification, and discrimination (Sniffin’ Sticks) behaviorally. Structural 3T MRI measured olfactory bulb volume, sulcus depth, and sulcus length. Sample sizes vary per analysis.

**Results:** Relative to TD, 22q11DS showed elevated sinonasal burden (p<.001), lower odor identification (p<.001), and lower odor discrimination (p<.001). PS showed elevated sinonasal burden (p<.001) and marginally lower odor identification (p=.06), but no discrimination deficit. Olfactory bulb volume was significantly smaller in 22q11DS versus TD (p=.007, r=.39); PS showed a non-significant trend (p=.07). Olfactory sulcus depth (p=.52) and length (p=.82) did not differ across groups.

**Conclusions:** 22q11DS is associated with elevated sinonasal burden, robust olfactory identification and discrimination deficits exceeding those of PS and reduced olfactory bulb volume. The co-occurrence of functional deficits and reduced bulb volume without sulcal anomalies may distinguish the 22q11DS olfactory phenotype from that of PS. These hypothesis-generating findings are consistent with partially dissociable neurodevelopmental mechanisms and support the utility of 22q11DS as a genetic model for psychosis risk.

## Introduction

The 22q11.2 deletion syndrome (22q11DS) is the most common chromosomal microdeletion in humans,(McDonald-McGinn et al., 2015) affecting approximately 1 in 4,000 live births and conferring the highest known genetic risk for schizophrenia-spectrum disorders outside of monogenic mutations. Approximately 25% of individuals with 22q11DS develop a psychotic disorder (Chow et al., 1999; Murphy et al., 1999), with clinical and neurobiological features that closely parallel those observed in schizophrenia (SZ) in the general population (Tang et al., 2017; Tang et al., 2016). Critically, 1–2% of individuals with SZ carry a 22q11.2 deletion (Bassett et al., 2010), underscoring the etiological relevance of this neurogenetic condition for understanding the broader psychosis spectrum (PS). 22q11DS thus offers a unique window into the neurodevelopmental mechanisms underlying psychosis risk (Gothelf et al., 2007; Tang and Gur, 2017). One neurobiological domain particularly well-suited to probing the neurodevelopmental origins of psychosis risk in 22q11DS is the olfactory system.

Olfactory dysfunction is a consistent and well-replicated finding in SZ. Meta-analytic evidence indicates medium-to-large deficits in odor identification (naming or recognizing an odor), odor discrimination (telling odors apart), and odor detection threshold (the lowest perceptible concentration) in patients with SZ, with smaller and more heterogeneous deficits in youth at clinical high risk for psychosis (Moberg et al., 2014). Several features of the peripheral olfactory system make it a particularly promising entry point for understanding the neurodevelopmental neuropathology of psychosis (Sawa and Cascella, 2009). Morphological differentiation of the peripheral olfactory system begins very early in utero, as nasal cavities are fully sculpted by gestational week 11, and olfactory receptor neuron axons reach the olfactory bulb by the late first trimester (Farbman, 1991; Larsen, 2001). This developmental window overlaps precisely with the critical embryonic period during which environmental dysregulation is thought to confer heightened risk for psychosis (Mednick et al., 1988). Deficits in odor identification are correlated with the severity of negative symptoms in patients with psychosis (Corcoran et al., 2005; Kästner et al., 2013) and are predictive of transition to a psychotic disorder in individuals at clinical high risk (Brewer et al., 2003; Woodberry et al., 2010). Olfactory identification deficits also predict poorer long-term functional outcome (Lin et al., 2015), making them candidates for early biomarker development.

In patients with SZ, structural changes exist at all levels of the ascending olfactory system, including the nasal cavities, olfactory bulbs, and primary olfactory cortex in the medial temporal lobe (Turetsky et al., 2009b), supporting the hypothesis that olfactory abnormalities reflect an early and pervasive disruption of midline neurodevelopment (Kamath et al., 2014; Turetsky et al., 2009b; Turetsky et al., 2018). Specifically, olfactory bulb volume is reduced in patients with SZ (Turetsky et al., 2000) and in clinical high-risk subjects (Turetsky et al., 2018), that is, youth with subthreshold psychotic symptoms, with males showing bilateral olfactory bulb reductions approximately 36% smaller than TD males. Reduced olfactory bulb volume in clinical high-risk subjects correlates with greater negative symptom burden (Turetsky et al., 2018), further linking peripheral structural integrity to the clinical expression of psychosis risk. Depth of the olfactory sulcus, a structural marker of early embryonic olfactory system development, is also reduced in SZ (Turetsky et al., 2009a) and has been proposed as a surrogate marker of prenatal disturbance.

Recent work has established that behavioral olfactory deficits, including impaired odor identification, discrimination and detection, are prominent in 22q11DS. Tang et al. (Tang et al., 2018) first documented robust olfactory identification and discrimination impairments in 22q11DS relative to typically developing (TD) controls and demonstrated that poorer odor discrimination was associated with greater psychosis-spectrum symptomatology within the 22q11DS group. A subsequent meta-analytic review of the extant literature confirmed the magnitude and consistency of these deficits across studies, with large and homogeneous impairments observed across odor identification, discrimination, and detection threshold tasks (Moberg et al., 2020). Notably, individuals with 22q11DS who also exhibited psychosis-spectrum symptoms (22q11DS+) showed greater odor discrimination deficits relative to those without psychosis symptoms (22q11DS−) (Tang et al., 2018), suggesting that olfactory measures may help index psychosis risk in 22q11DS. Despite this convergent evidence, little work has combined behavioral and structural olfactory measures in 22q11DS or compared these profiles directly with PS. Understanding whether the olfactory phenotype of 22q11DS mirrors that of PS, or whether it is distinguished by additional structural features reflecting its genetic and neurodevelopmental origins is critical for leveraging 22q11DS as a model for psychosis biomarker discovery.

The current study addresses this gap by characterizing a comprehensive battery of olfactory measures, including sinonasal symptom burden, behavioral odor identification and discrimination, olfactory bulb volume, and olfactory sulcal morphology across three groups: individuals with 22q11DS, PS youth without a known genetic syndrome (with subthreshold to threshold psychotic symptoms), and TD. We hypothesized that 22q11DS would exhibit significant deficits in behavioral olfactory function relative to TD, and that these deficits would exceed those observed in PS. The latter expectation was based on three considerations. First, meta-analytic olfactory deficits in 22q11DS are large and homogenous (Cohen’s d of ∼1.1 relative to TD) (Moberg et al., 2020), whereas deficits in youth at clinical risk for psychosis are more modest and heterogenous (Moberg et al., 2014). Second, 22q11DS is accompanied by craniofacial and midline abnormalities and by broad cognitive impairment, each of which may affect olfactory dysfunction. Third, PS youth are, by definition, largely subthreshold and early in the course of illness, when olfactory deficits are typically less pronounced than in SZ. With respect to structural measures, we hypothesized that 22q11DS would show lower olfactory bulb volume and reduced olfactory sulcal depth relative to TD, consistent with structural evidence of early olfactory system disruption in psychosis-related conditions. Because olfactory bulb volume in PS youth has been reported to be male-predominant (Turetsky et al., 2018), because sex differences in olfactory performance are well documented (Doty et al., 1984), and because the association between odor discrimination and psychosis-spectrum symptoms in 22q11DS was previously most evident in males (Tang et al., 2018), we also conducted exploratory sex-stratified analyses.

## Methods

Study methods are briefly described below. Complete details are provided in the Supplemental Methods.

### Participants

The sample included 209 participants: 101 22q11DS, 56 PS, and 52 TD. Individuals with 22q11DS were enrolled through the CHOP 22q and You Center and had confirmed A–D deletions and were enrolled irrespective of psychiatric status. Of the 91 22q11DS participants with consensus diagnostic data, n=67 (73.6%) had a lifetime psychotic disorder or psychosis-risk diagnosis (22q11DS+), including 11 with threshold psychotic disorder (12%). PS participants were recruited through the Penn–CHOP Lifespan Brain Institute and met clinically validated psychosis-spectrum criteria (≥1 positive or ≥2 negative/disorganized symptoms rated ≥3 on the Scale of Prodromal Symptoms [SOPS]) with persistent symptoms. All PS participants were psychosis spectrum at the time of the study. Lifetime diagnoses included threshold psychotic disorder (n=12, two with affective psychosis) and psychosis-risk syndromes (n=44), while the remaining participants carried secondary lifetime diagnoses of bipolar disorder (n=2) and depressive disorder (n=8) (Supplemental Table 8). TD participants had no current or past psychotic disorder or substance dependence (except nicotine) and no active psychiatric diagnosis at time of the study. No PS or TD participant had a known genetic syndrome. However, PS and TD participants were not genotyped, and the PS group is therefore described as lacking a known genetic etiology. Individuals were only enrolled if they were free of acute or chronic respiratory illness.

Participants ranged in age from 5–60 years in 22q11DS, 14–32 years in PS, and 16–34 years in TD. Not all participants completed all measures due to COVID-19 disruptions, participant burden, or MRI contraindications. Available sample sizes are reported per analysis. Current nicotine use and psychotropic medication status by group are reported in Table 1. All participants provided written informed consent (or assent with parental consent), and procedures were approved by the University of Pennsylvania and CHOP Institutional Review Boards.

**Table 1.** Sample Characteristics by Diagnostic Group.

| Variable | 22q11DS<br>(n=101) | PS<br>(n=56) | TD<br>(n=52) | 22q vs PS | 22q vs<br>TD | PS vs<br>TD |
| --- | --- | --- | --- | --- | --- | --- |
| **Demographics** |  |  |  |  |  |  |
| Sex, Male, n (%) | 48 (47.5%) | 34 (60.7%) | 28 (53.8%) | $\chi^2=2.56$, p=.28 | | |
| Age, M (SD) | 20.1 (8.9) | 23.7 (3.5) | 23.0 (4.2) | **p<.01** | **p=.03** | p=.35 |
| Race, White, n (%) | 82 (81.2%) | 24 (42.9%) | 26 (50.0%) | $\chi^2=27.90$, p<.001 | | |
| Tobacco use (past 12 months) | 1 (1.6%) | 10 (19.6%) | 4 (8.2%) | **Fisher's exact test, p=0.004<sup>#</sup>** |  |  |
| Current psychotropic, n (%) | 57 (62.0%) | 24 (42.9%) | 4 (7.8%) | $\chi^2=39.27$, p<.001 | | |
| Current antipsychotics, n (%) | 13 (14.1%) | 8 (14.3%) | 0 (0.0%) | $\chi^2=8.09$, p=.02 | | |
| **Cognitive Screening** |  |  |  |  |  |  |
| TONI-IQ, M (SD) [n] | 80.3 (12.6) [72] | 101.9 (17.6) [54] | 105.0 (20.3) [51] | **p<.001** | **p<.001** | p=.41 |
| MoCA, M (SD) [n] | 21.1 (4.8) [64] | 26.0 (2.9) [50] | 27.6 (2.2) [46] | **p<.001** | **p<.001** | **p=.004** |
| CCI, M (SD) [n] | 35.7 (14.5) [73] | 33.8 (13.4) [54] | 23.9 (5.2) [51] | p=.45 | **p<.001** | **p<.001** |

**Note.** 22q11DS (22q11.2 deletion syndrome); PS = Psychosis Spectrum; TD = Typically Developing; TONI-IQ = Test of Nonverbal Intelligence IQ; MoCA = Montreal Cognitive Assessment (higher = better); CCI = Cognitive Complaints Index (higher = more complaints). Pairwise $p$ -values from independent-samples $t$ -tests (continuous) or chi-square tests (categorical). Medication exposure reflects current use, which was not available for all participants (92 22q11DS, 56 PS and 51 TD had available medication data). Tobacco use was not available for all participants (62 22q11DS, 51 PS and 49 TD had available tobacco data). Bold $p$ -values indicate statistical significance ( $p<.05$ ). M (SD) [n] format used for continuous variables; not all participants completed each measure. <sup>#</sup>Fisher's exact test used because expected cell counts were below 5.

### Clinical and Cognitive Assessment

Psychiatric diagnoses were established using structured interviews administered by trained assessors. Clinical evaluation included the Schedule for Affective Disorders and Schizophrenia for School-Age Children, Present and Lifetime Version (K-SADS) (Kaufman et al., 1997), the Computer Assisted Psychopathology Assessment (CAPA) (Calkins et al., 2014; Tang et al., 2014), and/or the Structured Clinical Interview for DSM Disorders (SCID) (First et al., 2002). All participants completed the Structured Interview for Prodromal Syndromes (SIPS v. 4.0) (McGlashan et al., 2001), with symptom severity rated using the SOPS (Miller et al., 1999). Collateral parent interviews were obtained for minors. Final diagnoses were determined via consensus case conference led by doctoral-level clinicians. Medication exposure (any psychotropic or antipsychotic) was coded from medication records, and tobacco use from self-report substance questionnaire. Nonverbal intellectual ability was assessed with the Test of Nonverbal Intelligence–Fourth Edition (TONI-4) (Brown, 2003). Global cognitive function and subjective cognitive complaints were screened using the Montreal Cognitive Assessment (MoCA) (Nasreddine et al., 2005) and Cognitive Complaints Index (CCI) (Wang et al., 2006).

### Behavioral Assessment of Olfactory Function

Olfactory testing was conducted during a single session. Prior to testing, participants completed the Sino-Nasal Outcome Test (SNOT) (Piccirillo et al., 2002), a validated 20-item clinical inventory assessing the burden of sinonasal symptoms including nasal obstruction, discharge, facial pain, and associated quality-of-life impairments.

### Olfactory Psychophysics

The 16-item Sniffin’ Sticks Odor Identification Test (SS-OIT) and the 16-item Sniffin’ Sticks Odor Discrimination Test (Burghart Medizintechnik, Wedel, Germany) (Kobal et al., 1996) were administered birhinally according to standard instructions. Total correct responses were recorded for each test.

### Neuroimaging Protocol

Structural MRI data were acquired during a 35-minute session using a research-dedicated 3T Siemens Prisma scanner. All structural outcomes were derived from manual measurements on T1- and T2-weighted images. Scan quality was quantified with Euler number from FreeSurfer (v7.1.1) reconstruction of the T1-weighted image (averaged across hemispheres), a validated index sensitive to in-scanner head motion.(Rosen et al., 2018) All images were also visually inspected prior to tracing by raters blind to diagnosis.

### Olfactory Bulb Volume MRI

High-resolution T2-weighted 3D turbo-spin echo images (0.8 mm isotropic resolution) were acquired for manual olfactory bulb delineation (Turetsky et al., 2018) (Figure 1). Bilateral bulb volumes were manually traced by two raters blind to diagnosis. Inter-rater reliability exceeded r = .85. The primary outcome was bilateral bulb volume (mm^3^) averaged across hemispheres and raters. Hemisphere-specific volumes were examined in sensitivity analyses.

**Figure 1.**
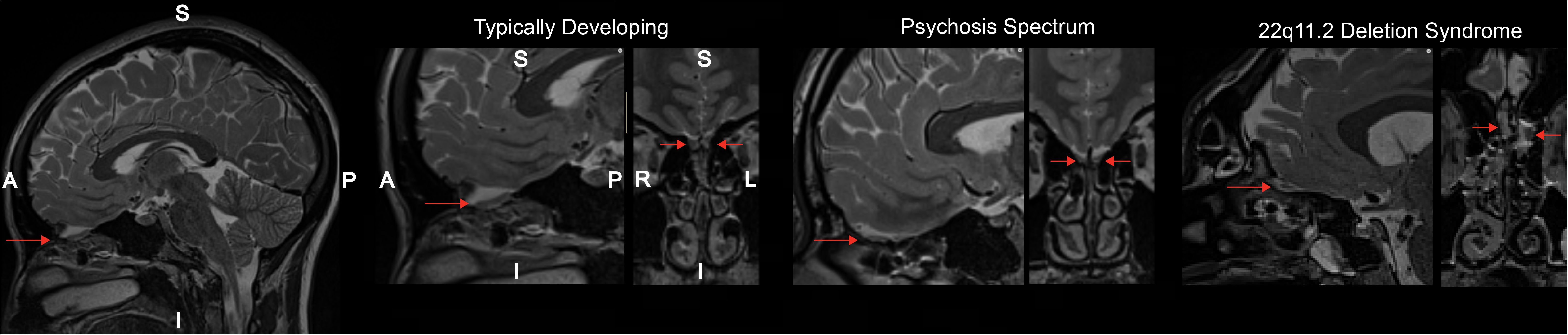
Representative T2-weighted MRI images of the olfactory bulb across diagnostic groups. The leftmost panel shows a full sagittal T2-weighted image illustrating the anatomical location of the olfactory bulb (red arrow) at the anterior base of the brain. The remaining panels display a 4x zoomed sagittal (left) and coronal (right) views of the olfactory bulb region for a representative participant from each diagnostic group: Typically Developing (TD), Psychosis Spectrum (PS), and 22q11.2 Deletion Syndrome (22q11DS). Red arrows indicate the olfactory bulb in each view. Orientation labels: A = anterior, P = posterior, S = superior, I = inferior, R = right, L = left.

### Olfactory Sulcal Morphology

Whole-brain T1-weighted MPRAGE images (0.96×0.96×1.0mm resolution) were used to measure olfactory sulcus depth and length. Scans were aligned to the AC–PC plane, and a standardized coronal reference slice was used for measurement. Sulcal depth was defined as the perpendicular distance from a tangent line spanning the gyrus rectus and medial orbital gyrus to the deepest point of the sulcus (Huart et al., 2011). Length was calculated as the average measurable extent across coronal slices. Measurements were performed by two raters blinded to group (CM, AA) using ImageJ (National Institutes of Health, Bethesda, MD), and values averaged across hemispheres and raters were used in primary analyses.

### Statistical Analysis

All statistical analyses were conducted in R (version 4.3) (R-Core-Team, 2012). Given that not all participants completed every measure, available sample sizes are reported for each analysis. Statistical significance was set at *p*<.05 (two-tailed) for all tests, and effect sizes are reported alongside all inferential statistics. Demographic and cognitive variables were compared using ANOVAs with Bonferroni-corrected pairwise t-tests for continuous variables and Pearson chi-square tests (Cramér’s V) for categorical variables.

Primary olfactory and sinonasal outcomes were compared across the three groups using Kruskal-Wallis tests due to unequal and, in some cases, small sample sizes. Where omnibus Kruskal-Wallis tests were significant, pairwise group comparisons were conducted using Mann-Whitney *U* tests with Bonferroni correction. The rank-biserial correlation (*r*) is reported as the effect size for all non-parametric pairwise comparisons. Exploratory age-partial (ρ) Spearman correlations were computed between all primary olfactory and sinonasal measures, both across the full sample and within each diagnostic group, and between these measures and the four SIPS subscale totals. Given the exploratory nature and number of correlations, uncorrected *p*-values are reported for these tests and all findings should be considered hypothesis-generating. To examine whether olfactory measures were moderated by sex, supplementary analyses stratified by sex were conducted repeating the above approaches. Sex-stratified analyses were exploratory and were not corrected for the additional comparisons. Because the groups differed in age, race and cognitive ability, several sensitivity analyses were conducted. First, all primary analyses were repeated after excluding two 22q11DS participants with ages substantially exceeding the remainder of the sample (ages 60 and 51 years; all other participants ≤ 38 years). Second, parametric ANOVAs were run for all primary group comparisons. Third, each primary outcome was modeled with least-squares regression on group, age (linear and quadratic) and sex with heteroscedasticity-consistent (HC) standard errors, and then additionally adjusted for race, nonverbal IQ, current tobacco use and for MRI outcomes, the Euler number (Supplemental Tables 6 and 11). Fourth, all nonparametric analyses were repeated after restricting all groups to the overlapping age range of 14 to 34 years (Supplemental Table 7). Fifth, primary comparisons were repeated after excluding 22q11DS+ participant and after excluding current tobacco users (Supplemental Tables 10 and 11).

## Results

### Participant Characteristics

Participant characteristics and statistics are shown in Table 1. Groups did not differ significantly in sex distribution. A significant overall group difference in age was observed (*F*(2, 206)= 5.92, *p*=.003). 22q11DS participants were on average younger than both PS and TD, reflecting the broader age range of the 22q11DS cohort. Groups also differed significantly in racial composition (χ^2^=27.90, *p* < .001), with the 22q11DS group comprising a higher proportion of White participants (81.2%) relative to PS (42.9%) and TD (50.0%).

### Cognitive and Screening Characteristics

Groups differed significantly on nonverbal intellectual ability, global cognitive screening, and subjective cognitive complaints (Table 1). 22q11DS scored significantly lower than PS and TD on all three measures, and PS scored below TD on the MoCA and CCI.

### Sinonasal Symptom Burden

Sinonasal symptom burden, assessed via the SNOT, differed significantly across groups (Kruskal-Wallis *H*=22.72, *p* < .001; Figure 2, left panel). Pairwise comparisons revealed that both 22q11DS and PS reported significantly greater sinonasal symptoms than TD, with moderate-to-large effect sizes (Supplemental Table 2). The 22q11DS and PS groups did not differ significantly from one another, suggesting a comparably elevated sinonasal symptom profile in both clinical groups relative to healthy controls. Sinonasal burden was elevated in both clinical groups across both sexes (Supplemental Results).

**Figure 2.**
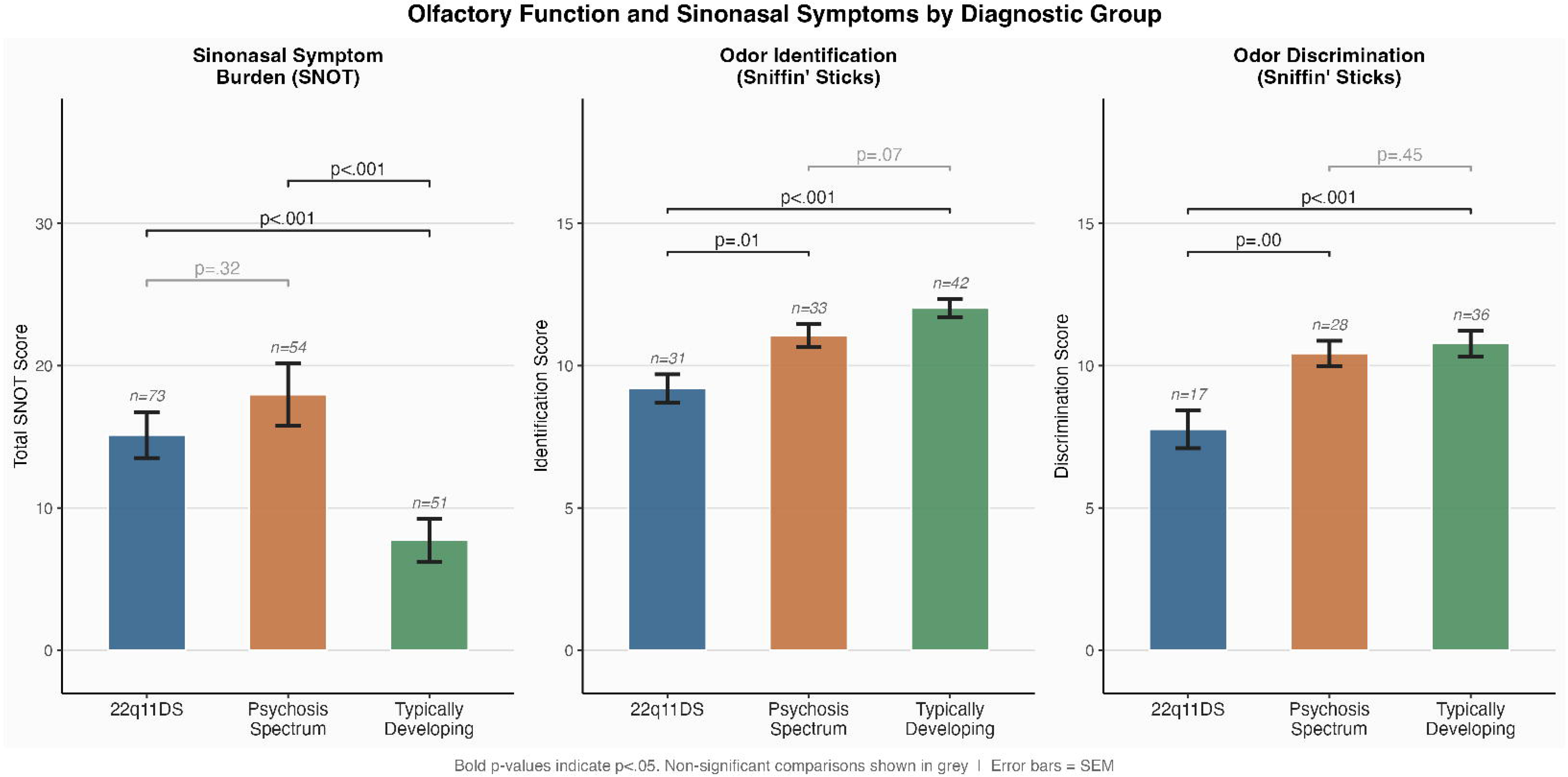
Olfactory function and sinonasal symptom burden by diagnostic group. Group means (± SEM) are shown for sinonasal symptom burden as measured by the Sino-Nasal Outcome Test total score (SNOT; left panel), odor identification score (center panel), and odor discrimination score (right panel) using the Sniffin’ Sticks battery, across 22q11.2 Deletion Syndrome (22q11DS), Psychosis Spectrum (PS), and Typically Developing (TD) groups. Sample sizes reflect participants with available data for each measure. Pairwise comparisons were conducted using Mann-Whitney U tests with Bonferroni correction (k = 3); uncorrected p-values are displayed on brackets. Bold p-values indicate p < .05 after Bonferroni correction; non-significant comparisons are shown in grey. Higher SNOT scores indicate greater sinonasal symptom burden; higher identification and discrimination scores indicate better olfactory performance. Error bars = SEM.

### Odor Identification

A significant overall group difference in odor identification performance was observed (*H* = 20.63, *p* < .001; Figure 2, middle panel). 22q11DS participants demonstrated markedly lower identification scores relative to both PS and TD. The PS and TD groups did not differ significantly on odor identification, though a trend in the expected direction was observed (Supplemental Table 2). The identification deficit in 22q11DS was of equivalent magnitude in males and females (Supplemental Results).

### Odor Discrimination

A significant overall group difference was also found for odor discrimination (*H*=13.21, *p*=.001; Figure 2, right panel). As with odor identification, 22q11DS participants performed significantly worse than both PS and TD. The PS and TD groups did not differ (Supplemental Table 2). Across both behavioral olfactory measures, effect sizes for the 22q11DS versus TD comparisons were large, indicating robust and clinically meaningful olfactory deficits in the 22q11DS group. The discrimination deficit in 22q11DS was directionally consistent across sexes (Supplemental Results).

### Olfactory Bulb Volume

A significant overall group difference in average bilateral olfactory bulb volume was observed (*H*=7.99, *p*=.018; Figure 3, left panel). Pairwise comparisons revealed that 22q11DS participants had significantly smaller olfactory bulb volumes relative to TD. The 22q11DS versus PS comparison did not reach statistical significance, nor did the PS versus TD comparison, though both trended in the expected direction (Supplemental Table 2). Hemisphere-specific sensitivity analyses revealed a consistent pattern, with 22q11DS showing significantly smaller volumes than TD for both left (*p*=.006) and right (*p*=.018) olfactory bulbs (Supplemental Figure 1).

**Figure 3.**
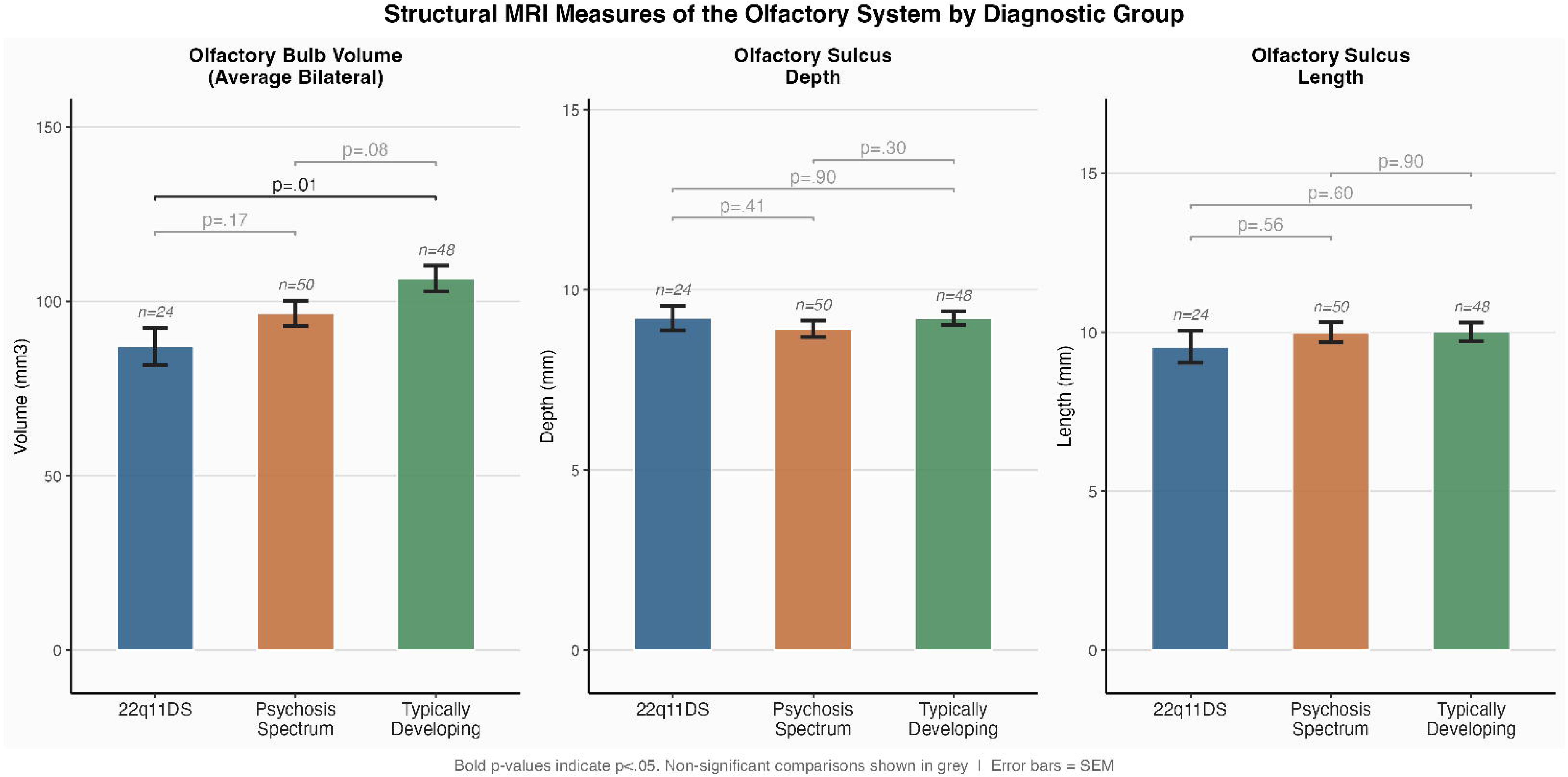
Structural MRI measures of the peripheral olfactory system by diagnostic group. Group means (± SEM) are shown for bilateral average olfactory bulb volume (mm^3^; left panel), olfactory sulcus depth (mm; center panel), and olfactory sulcus length (mm; right panel) across 22q11.2 Deletion Syndrome (22q11DS), Psychosis Spectrum (PS), and Typically Developing (TD) groups. Sample sizes reflect participants with available neuroimaging data for each measure. Pairwise comparisons were conducted using Mann-Whitney U tests with Bonferroni correction (k = 3); uncorrected p-values are displayed on brackets. Bold p-values indicate p < .05 after Bonferroni correction; non-significant comparisons are shown in grey. Error bars = SEM.

Among males, omnibus group differences were significant (Kruskal-Wallis *H*(2)=6.39, *p*=.04; Supplemental Figure 2), whereas group differences among females did not reach significance (*H*(2)=3.47, *p*=.18). Within males, both 22q11DS and PS showed trend-level reductions relative to TD that did not survive correction (Supplemental Results).

### Olfactory Sulcal Morphology

No significant group differences were observed for either olfactory sulcus depth (Figure 3, middle panel) or olfactory sulcus length (Figure 3, right panel). Complete details for all olfactory measures are provided in Supplemental Table 2.

### Sensitivity Analysis

Exclusion of the two oldest participants and parametric ANOVAs yielded results equivalent to the primary analyses. Scan quality did not differ across groups (H(2) = 1.77, p=0.41), was unrelated to MRI outcomes (all p>0.06), and did not alter group contrasts when included as a covariate (Supplemental Table 6, Supplemental Figure 3). In age- and sex-adjusted models, group effects were unchanged for sinonasal burden, odor identification (22q11DS vs TD), and odor discrimination (all corrected p<0.012). The 22q11DS vs TD bulb volume reduction was of equivalent magnitude, but fell short of corrected significance (corrected p=0.053), and the 22q11DS vs PS identification difference, while significant in analyses restricted to the overlapping age range of 14 to 34 years (p =0.03; Supplemental Table 7), was attenuated under parametric age adjustment (p=0.09). Group-by-age interactions were non-significant for all outcomes. Further adjustment for race and nonverbal IQ left sinonasal and behavioral effects intact but attenuated the bulb volume contrast (p=0.17; Supplemental Table 6).

22q11DS+ and 22q11DS- participants did not differ significantly on any olfactory measure, and the odor identification deficit remained large and significant relative to both TD and PS after excluding 22q11DS+ (Supplemental Table 10). Current psychotropic medication was more common in 22q11DS (62%) and PS (43%) than TD (10%). Current antipsychotic use (14% of each clinical group, no TD participants) was associated with higher sinonasal burden but with no other olfactory measure (Supplemental Table 9). Tobacco use was least common in 22q11DS (past 12-month use: 22q11DS = 1.6%; PS = 19.6%; TD = 8.2%; Fisher exact p = 0.004). Excluding current tobacco users left every significant effect intact and tobacco use was unrelated to any olfactory outcome in adjusted models (Supplemental Table 11).

### Associations amongst olfactory measures

Exploratory correlational analyses among olfactory measures are reported in full in the Supplemental Results. Briefly, odor identification and discrimination were significantly correlated across the full sample (ρ=.47, *p* < .001) and within 22q11DS (ρ=.57, *p*= .01). Olfactory bulb volume was positively associated with sulcus depth across groups (ρ=.30, *p<*.01), consistent with shared early embryonic origins, though behavioral performance and bulb volume were not robustly correlated at the individual level. Notably, SNOT was inversely associated with odor identification only in TD (ρ=−.40, *p*<.01), not in either clinical group.

### Associations Between Olfactory Measures and Clinical Symptoms

Exploratory correlational analyses between olfactory measures and SOPS subscales are reported in full in the Supplemental Results. Briefly, SNOT scores were significantly associated with all four SOPS subscales within 22q11DS, most strongly with general symptoms (ρ=.58, *p* < .001), and with general (p=0.01) and negative (p=0.03) symptoms within PS, but with no subscale in TD. Within 22q11DS, sulcus length was positively associated with positive symptom severity (age-partial ρ=.68, *p*<.01, *n*=16), a counterintuitive finding requiring replication given the small subsample, whereas in PS, sulcus length was negatively associated with negative symptoms (ρ=−.31, *p*=.04). Bulb volume was not significantly associated with any subscale (all *p*s > .17).

## Discussion

The present study provides a comprehensive characterization of olfactory and sinonasal phenotypes across three well-characterized groups, individuals with 22q11DS, PS youth without a known genetic syndrome, and typically developing controls, using both behavioral and structural measures of olfactory system integrity. Sinonasal symptom burden was elevated in both clinical groups and tracked psychosis-spectrum symptom severity. 22q11DS showed large odor identification and discrimination deficits relative to both PS and TD, and reduced olfactory bulb volume relative to TD, with exploratory analyses suggesting the bulb volume differences was more evident among males. Olfactory sulcus depth and length did not differ across groups. Taken together, these findings extend prior work by delineating a specific olfactory phenotype in 22q11DS that is characterized by pervasive functional deficits and peripheral structural volume reduction, but preserved sulcal morphology, a pattern that both overlaps with and diverges from what has been reported in SZ.

Both 22q11DS and PS groups reported significantly greater sinonasal symptom burden than TD, with medium-to-large effect sizes, and did not differ from one another. Sinonasal symptoms have received comparatively little attention in the psychosis literature, and their elevation in both clinical groups is noteworthy. Sinonasal inflammation and obstruction can impair olfactory receptor access and degrade psychophysical olfactory performance independent of central processing deficits (Kohler et al., 2001). The equivalent sinonasal symptom burden across 22q11DS and PS, combined with the substantially worse behavioral olfactory performance in 22q11DS, argues that sinonasal pathology alone cannot account for the magnitude of olfactory deficits in 22q11DS and that central olfactory processing impairments make an important independent contribution. Consistent with this, sinonasal burden was inversely associated with identification only within TD, the group with intact central olfactory processing. Beyond olfactory performance, SNOT scores were significantly associated with all four SIPS symptom subscales within 22q11DS and with general and negative symptoms within PS. Sinonasal burden was also higher among antipsychotic-treated participants in both clinical groups and descriptively higher in 22q11DS+ than 22q11DS-, so medication effects and general illness burden may both contribute to sinonasal symptom elevation. To our knowledge these are the first data directly associating sinonasal burden to dimensional psychiatric symptom severity in either population, and they may reflect shared inflammatory or dysautonomic mechanisms rather than simply a peripheral sensory comorbidity (Strous and Shoenfeld, 2006). Sinonasal pathology warrants further study as a potentially modifiable contributor to sensory dysfunction.

The large-magnitude impairments in odor identification and odor discrimination observed in 22q11DS replicate and extend prior findings from our group and others (Tang et al., 2018; Turetsky et al., 2009b), and are consistent with a large meta-analytic effect across olfactory psychophysical tasks in this syndrome (Moberg et al., 2020). The inclusion of a PS comparison group further indicates that the magnitude of deficit in 22q11DS exceeds that of PS, with PS individuals showing only trends toward poorer performance relative to TD on both measures. Deficits were of comparable magnitude in males and females. Although 22q11DS participants also had lower nonverbal IQ, the discrimination task minimizes verbal and executive demands and adjustment for nonverbal IQ did not eliminate the 22q11DS versus TD deficits in identification or discrimination, arguing against a purely cognitive explanation. We note, however, that statically adjusting for a core feature of the syndrome can itself produce biased estimates and that residual confounding cannot be excluded. The mechanisms underlying these deficits may involve dopaminergic dysregulation arising from *COMT* haploinsufficiency at 22q11.2, given the dense dopaminergic innervation of the olfactory bulb and primary olfactory cortex and prior evidence that *COMT* Val158Met genotype modulates olfactory performance in schizophrenia (Kamath et al., 2012). Structural airway abnormalities associated with velopharyngeal insufficiency (VPI) do not appear to be the a major contributor, as prior work demonstrated equivalent identification deficits in 22q11DS participants with and without VPI (Tang et al., 2018).

Olfactory bulb volume was significantly reduced in 22q11DS relative to TD, consistent with prior reports in SZ (Turetsky et al., 2000) and PS (Turetsky et al., 2018). Reduced volume likely reflects a combination of decreased afferent drive from the nasal epithelium, which is consistent with receptor-level dysfunction documented in SZ (Turetsky et al., 2009b), and intrinsic developmental anomalies of the bulb arising from disruption of olfactory placode development (Maynard et al., 2001). The bulb’s projections to perirhinal, entorhinal, and orbitofrontal cortices position it at the intersection of peripheral olfactory and broader psychosis-relevant neuropathology (Turetsky et al., 2009b; Turetsky et al., 2000), and the absence of individual-level correlations between bulb volume and behavioral performance within any group suggests these represent partially dissociable dimensions of olfactory system disruption rather than a unitary deficit.

The PS–TD bulb comparison did not reach significance, likely because the mixed-sex psychosis spectrum neuroimaging subsample attenuated an effect previously shown to be male-predominant in this group (Turetsky et al., 2018). Exploratory sex-stratified analyses were consistent with this interpretation, although they should be read cautiously. The omnibus group differences in bulb volume was significant among males but not females, pairwise comparisons among males did not survive correction, and PS females showed numerically larger bulb volumes than PS males, a pattern consistent with the male-predominant reduction previously reported in PS (Turetsky et al., 2018). By contrast, 22q11DS males and females showed comparably reduced volumes, raising the possibility that bulb volume reduction in 22q11DS is expressed more uniformly across sexes than in PS. This distinction is hypothesis-generating given the small number of females in the neuroimaging sample and requires replication in larger, sex-equivalent samples. Several other null findings warrant comment. The absence of a PS versus TD difference in odor discrimination, and the trend-level difference in identification, are consistent with meta-analytic evidence that olfactory deficits in youth at clinical risk are smaller and more heterogeneous than in SZ and may also reflect the relatively preserved functioning and early illness stage of the present PS sample, which consisted largely of individuals at clinical risk for psychosis. The null PS versus TD bulb volume comparison, in turn, likely reflects both the mixed-sex composition of the subsample and limited overall sample size.

Olfactory sulcus depth and length did not differ across any group, a null result that was consistent across sex and stands in contrast to prior evidence of sulcal depth reduction in SZ (Turetsky et al., 2009a). This dissociation of robust behavioral and bulb volume deficits in the context of preserved sulcal morphology is theoretically informative. Turetsky and colleagues (Turetsky et al., 2009b) have proposed that functional olfactory deficits and sulcal structural anomalies represent partially dissociable markers of distinct etiological pathways with the former indexing genetic vulnerability, the latter reflecting intrauterine environmental disruption. Consistent with this framework, familial SZ shows functional olfactory deficits in unaffected relatives without sulcal anomalies (Compton et al., 2007; Griffiths et al., 1998). The 22q11DS pattern is compatible with this framework. A high-penetrance deletion is accompanied by large functional and bulb volume deficits without the sulcal changes reported in SZ. We emphasize, however, that the present study did not measure prenatal or postnatal environmental exposures, and a null group difference in sulcal morphology cannot by itself rule out environmental contributions to the 22q11DS olfactory phenotype. Postnatal clinical factors that differ between 22q11DS and PS, including chronic sinonasal and middle-ear disease, recurrent infection, cardiac and palatal surgery, medication exposure and cognitive impairment, may also shape olfactory function and structure and were not fully accounted for in this study. The null sulcal finding should be interpreted cautiously given the modest 22q11DS neuroimaging sample, but individual-level co-variation between bulb volume and sulcal measures suggests sulcal morphology retains individual-level relevance.

The convergence of behavioral olfactory deficits, olfactory bulb volume reduction, and preserved sulcal morphology in 22q11DS constitutes a phenotypic profile that is partially distinct from PS and could, in principle, inform multivariate biomarker frameworks for distinguishing these two routes to psychosis risk. Behavioral deficits alone are unlikely to be useful for risk stratification within 22q11DS, as they appear broadly across the group. Indeed, identification deficits were equally present in 22q11DS+ and 22q11DS-individuals. Prior work suggests odor discrimination may more specifically index psychosis-spectrum symptom severity within 22q11DS, particularly in males (Schneider et al., 2014; Tang et al., 2018), and this warrants direct investigation in symptom-stratified samples. Whether combining behavioral and structural olfactory can help distinguish these two routes to psychosis risk remains an empirical question.

Several limitations of the current study should be noted. PS and TD participants were not genotyped. Given that 22q11.2 deletions are found in roughly 1-2% of the individuals with SZ, and likely fewer subthreshold PS, we would expect fewer than one undetected deletion carrier in the PS. Moreover, no PS or TD participant presented with features prompting genetic referral. Nonetheless, we cannot formally exclude an undetected deletion. The 22q11DS neuroimaging subsample was modest, potentially biased by the exclusion of more medically complex participants with MRI contraindications, which may limit generalizability of the null sulcal finding. The 22q11DS group spanned a considerably broader age range at both ends of the age spectrum than the PS and TD groups, which introduces age-related heterogeneity into olfactory performance data, as both identification and discrimination show well-documented age-related decline (Doty et al., 1984). However, exclusion of the two oldest 22q11DS participants, covariate adjustment for age and sex, and restriction to the overlapping age range yielded an equivalent pattern of results, although the corrected bulb volume contrast was marginal in adjusted models. Covariate-adjusted sensitivity analyses including race and nonverbal IQ preserved the sinonasal and behavioral findings, though the bulb volume contrast attenuated. More broadly, the groups may also differ in other key factors, but only a limited set of covariates could be collected and adjusted for in the present study. Residual confounding by comorbid diagnoses and the medical history of 22q11DS cannot be excluded. Additionally, tobacco data was collected as self-report and was unavailable for 39 22q11DS participants, but missing data were largely among younger children who were unlikely to be regular smokers. Finally, olfactory data were collected during the COVID-19 pandemic, a period during which SARS-CoV-2 infection is well-documented to cause olfactory dysfunction (Whitcroft and Hummel, 2020). Although participants were excluded if presenting with acute respiratory illness, no formal COVID-19 testing was conducted, and undetected prior or subclinical infection cannot be ruled out as a potential contributor to olfactory performance variability in the current sample.

This study provides the most comprehensive characterization to date of the olfactory phenotype in 22q11DS, integrating sinonasal, behavioral, and structural neuroimaging measures and benchmarking the 22q11DS profile directly against PS and TD. magnitude functional olfactory deficits and reduced olfactory bulb volume, which exceeded those of PS and did not differ by sex, converge with preserved sulcal morphology in a pattern that is consistent with, but does not establish, a predominantly genetic contribution to olfactory system disruption in 22q11DS. Because we did not measure prenatal and postnatal environmental exposures, we cannot exclude environmental and clinical contributions, and the findings should be regarded as hypothesis-generating. These findings nonetheless support 22q11DS as a neurodevelopmental model for psychosis and motivate multi-modal olfactory phenotyping as a candidate biomarker strategy to be tested in larger, independent samples.

## Supporting information

Supplemental Methods and Results

## Acknowledgements

We thank the patients and families who participated in this study.

## Availability of data and materials

The data generated and/or analyzed during the current study is available. Please contact Dr. David Roalf with questions and considerations for data sharing upon reasonable requests. Analysis code is available via github repository and shared upon request.

## Author Information

### Contributors Statement

All authors contributed to the writing, editing, and approval of this manuscript. DRR, PJM and BIT conceptualized and designed the research. DMM, MEC, TBC, PF, RSG, DEM, AO, LR, OT, EHZ, BSE, KR & REG prepared, assisted with and/or collected data. DRR, AA, CM & JES performed aspects of the MRI analysis. DRR & KR performed the statistical analyses. DRR wrote the first draft of the manuscript. All authors critically reviewed the manuscript’s content and approved the final version for publication.

### Corresponding author

Correspondence to Dr. David Roalf

## Ethics declarations

### Ethics approval and consent to participate

Participants provided informed consent/assent and the study procedures were approved by the Institutional Review Boards at the Children’s Hospital of Philadelphia and the University of Pennsylvania. Participants’ privacy and confidentiality were ensured at every stage of the study. We confirm that all methods were performed in accordance with the relevant guidelines and regulations.

### Consent for publication

Not applicable

