## Supplemental Methods and Results for "Olfactory Deficits and Structural Anomalies of the Peripheral Olfactory System in 22q11.2 Deletion Syndrome"

***Clinical and Cognitive Assessment***

Participants underwent a comprehensive clinical assessment to permit psychiatric diagnosis and characterize psychosis-spectrum symptomatology. Trained assessors administered a computerized version of the Schedule for Affective Disorders and Schizophrenia for School-Age Children – Present and Lifetime Version (K-SADS-PL)^1^ to assess mood, anxiety, psychotic, prodromal, ADHD, and substance use disorders. Adults were assessed using the Structured Clinical Interview for DSM-IV-TR Axis I Disorders (SCID-I/P)^2^. The Structured Interview for Prodromal Syndromes (SIPS v4.0)^3,4^ was administered to all participants to assess psychosis-risk symptoms (Supplemental Table 1). Symptom severity was rated using the Scale of Prodromal Symptoms (SOPS).^4^ A parent or guardian collateral interview was conducted for minor participants. Consensus diagnoses were established at clinical case conference led by at least two doctoral-level clinicians, integrating all available assessment data.

Nonverbal intellectual ability was assessed with the Test of Nonverbal Intelligence, Fourth Edition (TONI-4),^5^ a language-free measure of abstract reasoning and intellectual ability appropriate for individuals ages 6 through 89 years. A brief cognitive screening was performed using the Montreal Cognitive Assessment (MoCA),^6^ a validated brief measure of global cognitive functioning. Subjective cognitive complaints were assessed using the Cognitive Complaints Index (CCI),^7^ a self-report questionnaire assessing perceived difficulties in memory and attention in everyday functioning.

***Odor Identification.*** Odor identification was assessed using the Sniffin’ Sticks Odor Identification Test (SS-OIT; Burghart Medizintechnik, Wedel, Germany),^8^ a commercially available, standardized test kit with well-established reliability and validity across clinical and research populations. Sixteen odorants (Blue version) were presented sequentially via felt-tipped pen dispensers. For each item, participants were first asked to freely name the odor and were then presented with four response options from which to select the correct identification. The total number of correctly identified odors (maximum = 16) served as the primary identification outcome.

***Odor Discrimination.*** Odor discrimination was assessed using the Sniffin’ Sticks Odor Discrimination Test,^8^ in which participants were presented with 16 sets of three odorants. Within each triplet, two odors were identical and one was different; participants were asked to identify the deviant stimulus. This paradigm eliminates potential confounds associated with verbal labeling, cognitive demands of identification, or response facilitation by verbal choice alternatives, providing a purer measure of peripheral and early-stage olfactory processing. The total number of correctly identified deviant odors (maximum = 16) served as the primary discrimination outcome.

***Olfactory Bulb Volume Acquisition.*** High-resolution images of the olfactory bulbs were acquired using a T2-weighted three-dimensional multi-slice turbo-spin echo sequence (TR = 9000 ms; TE = 63 ms; bandwidth = 190 Hz/pixel; FOV = 256 × 256 mm; 60 slices; flip angle = 180°; effective voxel resolution = 0.8 × 0.8 × 0.8 mm; scan time ≈ 6 minutes). This sequence has been used in prior studies of olfactory bulb morphometry in psychosis from our laboratory^9,10^ and provides sufficient resolution for reliable manual delineation of bulb boundaries. Bilateral olfactory bulb volumes were determined by two trained raters using manual tracing of the bulb contour across multiple slices in the sagittal plane. All tracers were trained to criterion and blind to diagnostic group assignment. Inter-rater reliability of manual tracing exceeded *r* = .85 in our laboratory. The primary outcome measure was the average bilateral olfactory bulb volume (mm³). Left and right hemisphere volumes are reported separately as sensitivity analyses (Supplemental Figure 1).

***Olfactory Sulcal Morphology.*** Measurements of olfactory sulcus depth and length were obtained from whole-brain T1-weighted MPRAGE images (TR = 1680 ms; TE = 4.67 ms; FOV = 135 × 240 mm; matrix = 180 × 240; effective voxel resolution = 0.96 × 0.96 × 1.0 mm). Prior to measurement, each participant’s scan was aligned in three dimensions orthogonal to the AC-PC line. A standard coronal reference slice was selected as the plane of the posterior tangent through the eyeballs (PPTE), defined as the most caudal coronal slice passing through both eyeballs. A straight line was drawn tangent to the surface of the gyrus rectus and the medial orbital gyrus. Olfactory sulcus depth was defined as the perpendicular distance from this tangent line to the deepest point of the sulcus, measured using a single voxel-width line.^11^ Olfactory sulcus length was calculated as the average linear extent of the sulcus across all measurable coronal slices. All measurements were made by two expert raters (CM, AA) using ImageJ (National Institutes of Health, Bethesda, MD; https://imagej.nih.gov/), blind to diagnostic group assignment. Left and right hemisphere measurements were averaged to yield bilateral means for the primary analyses.

Diagnosis, Medication, and Tobacco Data. For 22q11DS, lifetime psychiatric diagnoses were curated from consensus diagnosis records (available for 195 of 209 participants). Every DSM Axis I code across a participant’s records was classified and the highest-priority category in a fixed diagnosis was taken as the primary diagnosis. PS classifications were verified against the most recent clinical assessment. Medication exposure was coded from medication collection records by drug name. Any psychotropic was defined as an antipsychotic, antidepressant, ADHD medication, anxiolytic or mood stabilizer, and antipsychotic exposure was coded separately, each as documented current use and as lifetime documented use. Does was not readily available. Tobacco use was drawn from self-report substance inventory, with ever-use and past-12-month use coded from the primary tobacco question. Availability and group comparisons for these variable are reported in Supplement Tables 8, 9, and 11.

**Supplemental Results**

**Supplemental Table 1. Descriptive Statistics for SOPS Symptom Subscales by Diagnostic Group**

|  | **22q11DS** | | | | **Psychosis Spectrum** | | | | **Typically Developing** | | | |
| --- | --- | --- | --- | --- | --- | --- | --- | --- | --- | --- | --- | --- |
| **SIPS Subscale** | **M** | **SD** | **n** | **Range** | **M** | **SD** | **n** | **Range** | **M** | **SD** | **n** | **Range** |
| Positive | 5.07 | 4.66 | 73 | 0–22 | 8.69 | 6.71 | 52 | 0–27 | 1.57 | 1.94 | 51 | 0–8 |
| Negative | 9.60 | 6.18 | 73 | 0–31 | 7.90 | 4.26 | 52 | 0–18 | 1.94 | 2.77 | 51 | 0–12 |
| Disorganized | 4.79 | 4.06 | 73 | 0–18 | 4.37 | 3.07 | 52 | 0–14 | 0.63 | 1.31 | 51 | 0–8 |
| General | 5.32 | 4.65 | 73 | 0–24 | 5.50 | 3.56 | 52 | 0–13 | 1.33 | 1.93 | 51 | 0–7 |

*Note. M = mean; SD = standard deviation; n = valid sample size for each measure;*

**Supplemental Table 2. Descriptive Statistics and Group Comparisons for Olfactory and Sinonasal Measures**

|  | **22q11DS** | | **Psychosis Spectrum** | | **Typically Developing** | | **Omnibus** | **22q11DS vs PS** | | **22q11DS vs TD** | | **PS vs TD** | |
| --- | --- | --- | --- | --- | --- | --- | --- | --- | --- | --- | --- | --- | --- |
| **Measure** | **M (SD)** | **n** | **M (SD)** | **n** | **M (SD)** | **n** | **H (df = 2), p** | **r** | **p** | **r** | **p** | **r** | **p** |
| SNOT | 15 (14) | 73 | 18 (16) | 54 | 8 (11) | 51 | 22.72, < .001 | .10 | .323 | -.40 | **< .001*** | -.51 | **< .001*** |
| Odor Identification | 9.19 (2.80) | 31 | 11.06 (2.30) | 33 | 12.02 (2.04) | 42 | 20.63, < .001 | .40 | **.006*** | .61 | **< .001*** | .25 | .067 |
| Odor Discrimination | 7.76 (2.75) | 17 | 10.43 (2.36) | 28 | 10.78 (2.73) | 36 | 13.21, .001 | .52 | **.004*** | .59 | **< .001*** | .11 | .448 |
| Bulb Volume (mm³) | 87.10 (26.24) | 24 | 96.61 (25.45) | 50 | 106.58 (25.12) | 48 | 7.99, .018 | .20 | .169 | .39 | **.007*** | .21 | .077 |
| Sulcus Depth (mm) | 9.22 (1.68) | 24 | 8.92 (1.60) | 50 | 9.21 (1.27) | 48 | 1.31, .521 | -.12 | .409 | -.02 | .895 | .12 | .298 |
| Sulcus Length (mm) | 9.54 (2.46) | 24 | 9.99 (2.24) | 50 | 10.01 (2.07) | 48 | 0.39, .824 | .08 | .560 | .08 | .603 | -.01 | .901 |

*Note. M = mean; SD = standard deviation; n = valid sample size for each measure; H = Kruskal-Wallis test statistic (df = 2); r = rank-biserial correlation (effect size); p = uncorrected p-value from Mann-Whitney U test. * p-value remains significant after Bonferroni correction (k = 3 comparisons per measure; p_adj < .05). SNOT = Sinonasal Outcome Test-20 total score; Bulb Volume = average bilateral olfactory bulb volume (mm³).*

**Hemisphere-Stratified Olfactory Bulb Analysis**

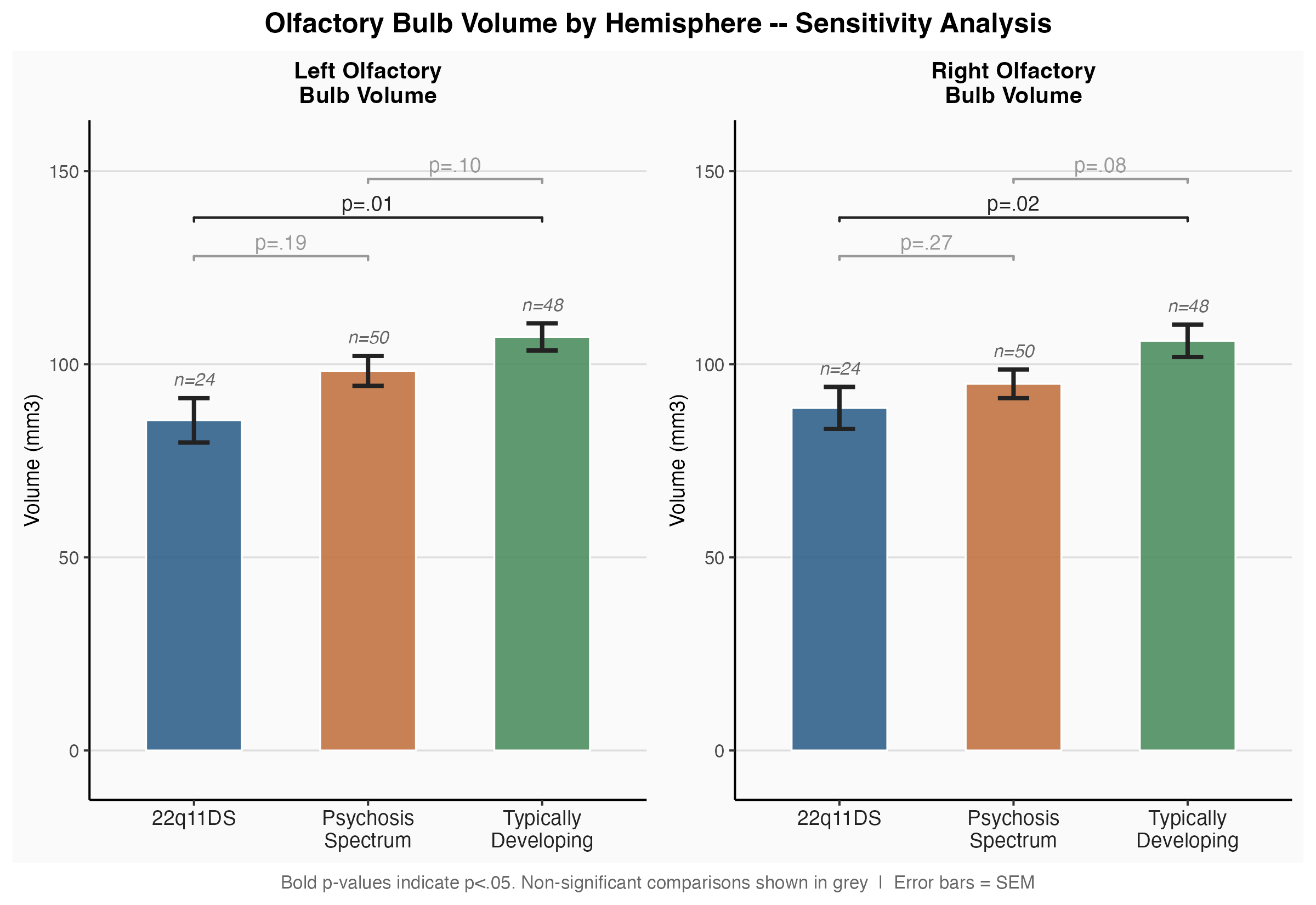

**Supplemental Figure 1. Olfactory bulb volume by hemisphere and diagnostic group.** Group means (± SEM) are shown for left (left panel) and right (right panel) olfactory bulb volume (mm³) across 22q11.2 Deletion Syndrome (22q11DS), Psychosis Spectrum (PS), and Typically Developing (TD) groups. Pairwise comparisons were conducted using Mann-Whitney U tests with Bonferroni correction (k = 3); uncorrected p-values are displayed on brackets. Bold p-values indicate p < .05 after Bonferroni correction; non-significant comparisons are shown in grey. Error bars = SEM

**Sex-Stratified Analyses**

*Sinonasal Symptom Burden (SNOT).* Sex-stratified analyses were exploratory and were not corrected for the additional comparisons they entail. Findings should be considered hypothesis-generating. Group differences on the SNOT were significant in both males (H = 7.67, p = .02) and females (H = 18.66, p < .01). In both sexes, 22q11DS and PS showed elevated SNOT scores relative to TD and did not differ from each other. PS females vs. TD females (p_adj < .01, r = .68) and 22q11DS females vs. TD females (p_adj < .01, r = .56) showed large effect sizes across comparisons. In males, PS showed higher scores than TD (p_adj = .03, r = .39). 22q11DS males showed trend level higher scores than TD (p_adj = .07, r = .33).

*Odor Identification.* Group differences were significant in both males (H = 9.68, p < .01) and females (H = 10.99, p < .01). In both sexes, the pattern was identical: 22q11DS shows significant deficits relative to TD (males: p_adj < .01, r = .60; females: p_adj < .01, r = .61), while PS did not differ significantly from TD after correction (males: p_adj = .11; females: p_adj = .17). 22q11DS vs. PS did not differ for either sex.

*Odor Discrimination.* Group differences were significant in both males (H = 8.57, p = .01) and females (H = 7.61, p = .02), but the pairwise pattern diverges by sex. In males, 22q11DS was significantly worse than both PS (p_adj = .04, r = .58) and TD (p_adj = .01, r = .67), with PS and TD performing similarly. In females, 22q11DS showed trend level lower scores than TD (p_adj = .10, r = .54), and PS vs. TD was also non-significant (p_adj = .12, r = .50) even though both effects were medium-to-large likely due to small female cell sizes (22q11DS n = 8, PS n = 9).

*Olfactory Bulb Volume.* Among males, omnibus group differences were significant (H(2) = 6.39, p = .04), whereas group differences among females did not reach significance (H(2) = 3.47, p = .18). Within males, pairwise comparisons revealed a trend toward reduced bulb volume in 22q11DS males relative to TD males (p = .04, r = .43) and a similar trend for PS males relative to TD males (p = .03, r = .34), neither of which survived Bonferroni correction. 22q11DS and PS did not differ (p = 0.50). Notably, the directional pattern of sex differences within each clinical group diverged meaningfully: in 22q11DS, males and females showed comparably reduced bulb volumes (M = 83.3 vs. 90.3 mm³, Δ = 7.0 mm³), whereas in PS, females showed numerically larger bulb volumes than males (M = 106.1 vs. 91.3 mm³, Δ = 14.7 mm³). This pattern is consistent with prior reports of male-predominant bulb volume reductions in idiopathic psychosis spectrum samples. These findings suggest that bulb volume reduction in 22q11DS may be sex-nonspecific, in contrast to the more sexually dimorphic pattern observed in PS.

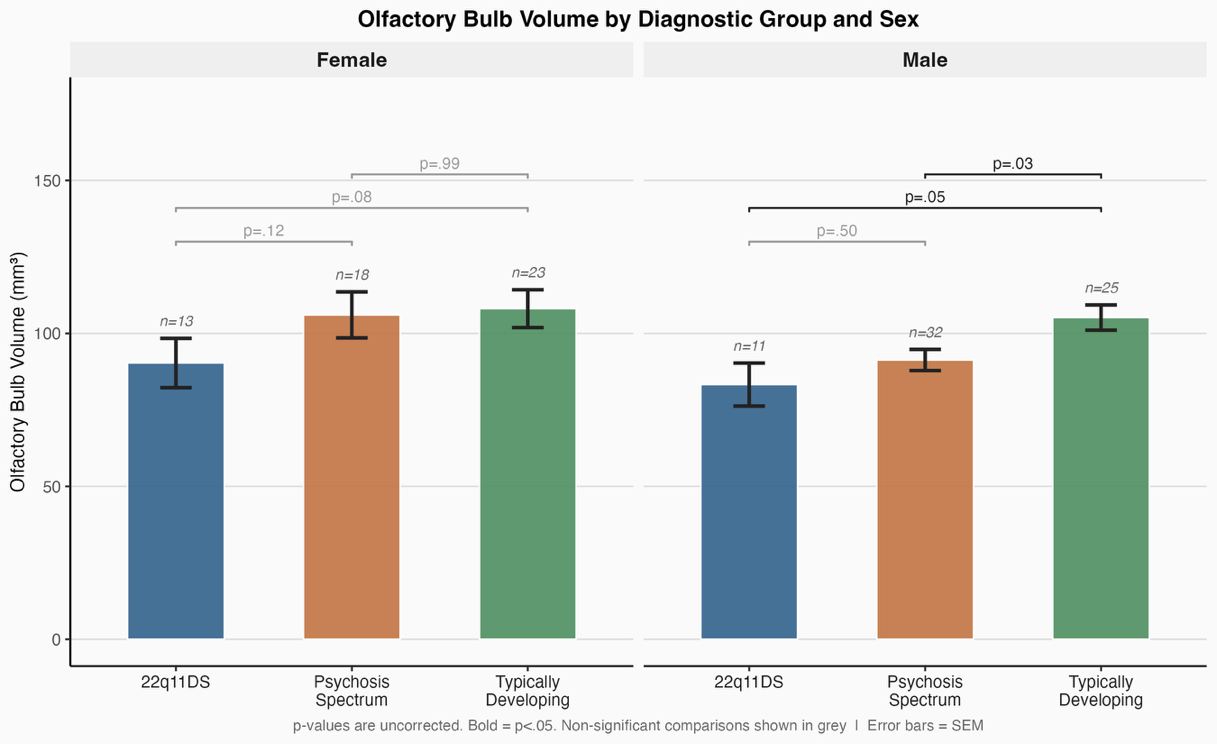

**Supplemental Figure 2 Olfactory bulb volume by diagnostic group and sex.** Group means (± SEM) for bilateral average olfactory bulb volume (mm³) are shown separately for females (left panel; n = 54) and males (right panel; n = 68) across 22q11.2 Deletion Syndrome (22q11DS), Psychosis Spectrum (PS), and Typically Developing (TD) groups. Pairwise comparisons were conducted using Mann-Whitney U tests with Bonferroni correction (k = 3); uncorrected p-values are displayed on brackets. Bold p-values indicate p < .05 (uncorrected); non-significant comparisons are shown in grey. Error bars = SEM.

*Olfactory Sulcus Depth.* Sulcus depth was non-significant in both males (H = 2.35, p = .31) and females (H = 3.24, p = .20). All pairwise comparisons were non-significant in both sexes (all p_adj ≥ .34).

*Olfactory Sulcus Length.* Sulcus length was non-significant in both males (H = 1.94, p = .38) and females (H = 1.09, p = .58). All pairwise comparisons were non-significant (all p_adj = 1.00).

**Associations Among Olfactory Measures**

Spearman rank correlations were computed between all primary olfactory measures both across the full sample and within each diagnostic group. Given the exploratory nature of these analyses and the multiple comparisons involved, findings are interpreted conservatively and descriptively; uncorrected p-values are reported alongside effect sizes to facilitate evaluation of magnitude. Complete correlation matrices are presented in Supplemental Table 3.

*Odor Identification and Odor Discrimination.* The strongest and most consistent association was between odor identification and odor discrimination scores (overall: ρ = .47, p < .01, n = 79; age-partial ρ = .47, p < .01). This association was significant within 22q11DS (ρ = .57, p = .02, n = 17; age-partial ρ = .57, p = .02) and TD (ρ = .49, p < .01, n = 35; age-partial ρ = .49, p < .01), and trended in the same direction within PS (ρ = .28, p = .16, n = 27), where power was more limited. The consistency of this association across diagnostic groups, including within 22q11DS despite that group's substantially lower mean performance on both tasks, indicates that individual variation in one olfactory behavioral domain reliably tracks variation in the other, and argues against floor effects dominating performance within 22q11DS.

*Sinonasal Symptom Burden and Behavioral Olfactory Performance.* SNOT scores were not significantly associated with odor identification or odor discrimination in the overall sample (identification: ρ = −.16, p = .13, n = 93; discrimination: ρ = −.06, p = .64, n = 74), nor within 22q11DS or PS (all ps > .25). A notable exception was observed within the TD group, where greater sinonasal symptom burden was significantly associated with poorer odor identification performance (ρ = −.40, p < .01, n = 42). No analogous association was present in either clinical group. This dissociation is consistent with the interpretation that olfactory behavioral deficits in 22q11DS and PS are driven predominantly by central olfactory processing impairments rather than by peripheral sinonasal obstruction: in individuals with intact central olfactory processing (TD), nasal symptom burden meaningfully constrains identification performance, but this peripheral pathway is obscured in clinical groups where central deficits dominate.

**Supplemental Table 3. Exploratory Correlations Among Primary Olfactory Measures by Diagnostic Group**

|  | **22q11DS** | | | **Psychosis Spectrum** | | | **Typically Developing** | | |
| --- | --- | --- | --- | --- | --- | --- | --- | --- | --- |
| **Pair** | **ρ** | **p** | **n** | **ρ** | **p** | **n** | **ρ** | **p** | **n** |
| SNOT × Odor ID | .12 | .62 | 19 | .21 | .26 | 32 | -.40 | **< .01** | 42 |
| SNOT × Odor Disc | .38 | .25 | 11 | .20 | .31 | 27 | -.23 | .17 | 36 |
| SNOT × Bulb Volume | -.33 | .21 | 16 | -.26 | .08 | 49 | -.02 | .87 | 47 |
| SNOT × Sulcus Depth | .03 | .92 | 16 | -.21 | .15 | 49 | .04 | .79 | 47 |
| SNOT × Sulcus Length | .32 | .23 | 16 | -.18 | .21 | 49 | .03 | .87 | 47 |
| Odor ID × Odor Disc | .57 | **.02** | 17 | .28 | .16 | 27 | .49 | **< .01** | 35 |
| Odor ID × Bulb Volume | .32 | .20 | 18 | .02 | .93 | 31 | -.02 | .91 | 38 |
| Odor ID × Sulcus Depth | .42 | .09 | 18 | .20 | .28 | 31 | .01 | .96 | 38 |
| Odor ID × Sulcus Length | .43 | .08 | 18 | .14 | .45 | 31 | .01 | .93 | 38 |
| Odor Disc × Bulb Volume | .37 | .21 | 13 | -.17 | .41 | 26 | .16 | .37 | 33 |
| Odor Disc × Sulcus Depth | .06 | .84 | 13 | .07 | .73 | 26 | .21 | .24 | 33 |
| Odor Disc × Sulcus Length | .13 | .68 | 13 | -.00 | 1.00 | 26 | .24 | .17 | 33 |
| Bulb Volume × Sulcus Depth | .27 | .21 | 24 | .34 | **.01** | 50 | .33 | **.02** | 48 |
| Bulb Volume × Sulcus Length | .06 | .78 | 24 | .34 | **.02** | 50 | .25 | .09 | 48 |
| Left Bulb Volume × Right Bulb Volume | .78 | **< .01** | 24 | .77 | **< .01** | 50 | .71 | **< .01** | 48 |

*Note. ρ = Spearman rank correlation; p = uncorrected p-value; n = valid sample size for each pair. Pairs with n < 8 within a group are not reported (—). Bold p-values indicate p < .05. All analyses are exploratory; uncorrected p-values are reported throughout.*

*Sinonasal Symptom Burden and Olfactory Bulb Volume.* Across the full sample, greater sinonasal symptom burden was associated with smaller bilateral olfactory bulb volume (ρ = −.21, p = .02, n = 112; age-partial ρ = −.21, p = .02), an association that persisted after controlling for age. This relationship was not significant within any individual group after correction, though a directionally consistent trend was observed within PS (ρ = −.26, p = .08, n = 49; age-partial ρ = −.27, p = .06). The overall association is plausible mechanistically: chronic sinonasal inflammation may reduce afferent olfactory receptor neuron input to the bulb, contributing to activity-dependent volume loss.

*Behavioral Olfactory Performance and Olfactory Bulb Volume.* Across the full sample, odor identification showed a trend toward a positive association with bilateral bulb volume (ρ = .20, p = .07, n = 87; age-partial ρ = .21, p = .05). Odor discrimination was not significantly associated with bulb volume overall (ρ = .14, p = .25, n = 72), and neither behavioral measure correlated significantly with bulb volume within any individual diagnostic group (all ps > .19 within 22q11DS, PS, and TD). The absence of robust behavioral–structural associations at the individual level, despite robust group-level differences in both domains within 22q11DS, suggests that olfactory bulb volume reduction and behavioral olfactory deficits reflect partially dissociable aspects of olfactory system disruption rather than a single underlying dimension.

*Olfactory Bulb Volume and Sulcal Morphology.* A consistent positive association was observed between olfactory bulb volume and olfactory sulcus depth across the full sample (ρ = .30, p < .01, n = 122; age-partial ρ = .31, p < .01) and within PS (ρ = .34, p = .02, n = 50; age-partial ρ = .34, p = .01) and TD (ρ = .33, p = .02, n = 48; age-partial ρ = .32, p = .03) separately. A parallel, somewhat weaker, association was observed between bulb volume and sulcus length overall (ρ = .24, p < .01, n = 122) and within PS (ρ = .34, p = .02, n = 50). Within 22q11DS, the bulb–sulcus depth association was directionally consistent but non-significant (ρ = .27, p = .21, n = 24), likely reflecting limited power in the neuroimaging subsample. These associations indicate that olfactory bulb volume and sulcal morphology co-vary at the individual level, even in groups where the mean sulcal depth does not differ from TD, consistent with their shared origins in early embryonic development of the olfactory placode and ventral forebrain. Importantly, this finding demonstrates that the group-level null result for sulcal morphology does not imply that sulcal depth is uninformative as an individual-differences measure, rather, it suggests that 22q11DS participants and controls are drawn from overlapping distributions of sulcal depth despite differing substantially in bulb volume and behavioral olfactory performance.

*Odor Identification and Sulcal Morphology within 22q11DS.* Within the 22q11DS group, odor identification scores showed positive trends with both sulcus depth (ρ = .42, p = .09, n = 18; age-partial ρ = .44, p = .07) and sulcus length (ρ = .43, p = .08, n = 18; age-partial ρ = .48, p = .04). The sulcus length association survived age correction and reached significance (age-partial ρ = .48, p = .04). No analogous associations between behavioral olfactory performance and sulcal morphology were observed within PS or TD (all ρ < .25, all ps > .17). Although these findings must be interpreted cautiously given the small within-group sample available for this analysis (n = 18), they suggest that within 22q11DS, individuals with greater sulcal extent, which likely reflects more intact early embryonic olfactory system development, tend to retain better olfactory identification performance. If replicated in larger samples, this within-group structural-functional relationship would provide further evidence that embryonic olfactory system development, as indexed by sulcal morphology, influences functional olfactory capacity even in the context of a shared genetic deletion.

*Bilateral Olfactory Bulb Symmetry.* Left and right olfactory bulb volumes were highly correlated across all groups (overall: ρ = .74, p < .01, n = 122; 22q11DS: ρ = .78, p < .01, n = 24; PS: ρ = .77, p < .01, n = 50; TD: ρ = .71, p < .01, n = 48), confirming high bilateral symmetry and supporting the use of the bilateral average as the primary structural outcome.

**Associations Between Olfactory Measures and SIPS Clinical Symptom Subscales**

Clinical symptom data from the Structured Interview for Prodromal Syndromes (SIPS v 4.0) were available for 73 of 101 22q11DS participants, 52 of 56 PS participants, and 51 of 52 TD participants. Scale of Prodromal Symptoms (SOPS) subscales examined were positive (P), negative (N), disorganized (D), and general (G) symptom totals. Descriptive statistics for SOPS subscales by group are presented in Supplemental Table 1. All correlations (Supplemental Table 4) are Spearman's ρ with age partialled out. Given the exploratory nature of these analyses, uncorrected p-values are reported alongside effect sizes. Findings are interpreted descriptively and should be considered hypothesis-generating.

*Sinonasal Symptom Burden and SIPS Subscales.* Within 22q11DS (n = 58), SNOT was significantly associated with all four subscales: general (ρ = .58, p < .01), disorganized (ρ = .40, p < .01), positive (ρ = .29, p = .03), and negative symptoms (ρ = .27, p = .04). Within PS (n = 50), significant associations were observed for general (ρ = .35, p = .01) and negative symptoms (ρ = .30, p = .03), with a trend for disorganized symptoms (ρ = .24, p = .09). In TD (n = 51), SNOT was not significantly associated with any SIPS subscale (positive: ρ = .13, p = .37; negative: ρ = .25, p = .08; disorganized: ρ = −.07, p = .64; general: ρ = .11, p = .43).

*Behavioral Olfactory Performance and SOPS Subscales.* Odor identification was not significantly associated with any SOPS subscale in either 22q11DS (all ps > .09) or PS (all ps > .13). Within 22q11DS, a trend was observed between odor identification and disorganized symptoms (age-partial ρ = .40, p = .09, n = 20). Odor discrimination showed significant positive associations with positive symptoms (ρ = .49, p = .01, n = 26) and general symptoms (ρ = .57, p < .01, n = 26) within PS. No significant associations between odor discrimination and SOPS subscales were observed within 22q11DS (all ps > .29, n = 14). The positive direction of the PS odor discrimination × symptom associations, indicating that higher discrimination scores co-vary with greater symptom severity, is counterintuitive and should be interpreted cautiously. It may reflect confounding with illness duration, medication status, or restricted range of discrimination scores in this sample.

*Olfactory Bulb Volume and SOPS Subscales.* Olfactory bulb volume was not significantly associated with any SIPS subscale in either 22q11DS (all ps > .44, n = 16), or PS (all ps > .17, n = 46).

*Olfactory Sulcal Morphology and SOPS Subscales.* Within 22q11DS, olfactory sulcus length showed a significant positive association with positive symptom severity (age-partial ρ = .68, p < .01, n = 16), with a parallel trend for sulcus depth (ρ = .49, p = .05, n = 16). No other significant associations between sulcal morphology and SOPS subscales were observed within 22q11DS (all remaining ps > .13). Within PS, sulcus length was significantly negatively associated with negative symptoms (ρ = −.31, p = .04, n = 46), and sulcus depth showed a trend in the same direction (ρ = −.28, p = .06, n = 46). All remaining PS sulcal × symptom associations were non-significant (all ps > .22). Complete correlation matrices for all olfactory × symptom pairs, within each group are presented in Supplemental Table 4.

**Supplemental Table 4. Correlations Between Olfactory Measures and SOPS Symptom Subscales**

|  |  | **22q11DS** | | | **Psychosis Spectrum** | | | **Typically Developing** | | |
| --- | --- | --- | --- | --- | --- | --- | --- | --- | --- | --- |
| **Olfactory Measure** | **SIPS Subscale** | **ρ** | **p** | **n** | **ρ** | **p** | **n** | **ρ** | **p** | **n** |
| SNOT | Positive | .29 | **.03** | 58 | .07 | .64 | 50 | .12 | .42 | 51 |
| SNOT | Negative | .27 | **.04** | 58 | .30 | **.03** | 50 | .25 | .08 | 51 |
| SNOT | Disorganized | .40 | **< .01** | 58 | .24 | .09 | 50 | -.08 | .58 | 51 |
| SNOT | General | .58 | **< .01** | 58 | .35 | **.01** | 50 | .12 | .41 | 51 |
| Odor ID | Positive | .24 | .30 | 20 | .06 | .76 | 31 | -.34 | **.03** | 42 |
| Odor ID | Negative | .20 | .40 | 20 | .06 | .74 | 31 | -.31 | **.04** | 42 |
| Odor ID | Disorganized | .39 | .09 | 20 | .06 | .75 | 31 | -.08 | .61 | 42 |
| Odor ID | General | .14 | .56 | 20 | .28 | .13 | 31 | -.29 | .06 | 42 |
| Odor Disc | Positive | .10 | .73 | 14 | .48 | **.01** | 26 | .03 | .88 | 36 |
| Odor Disc | Negative | -.30 | .29 | 14 | .25 | .21 | 26 | .11 | .53 | 36 |
| Odor Disc | Disorganized | -.00 | .99 | 14 | .23 | .27 | 26 | .06 | .71 | 36 |
| Odor Disc | General | .08 | .79 | 14 | .57 | **< .01** | 26 | -.05 | .79 | 36 |
| Bulb Volume | Positive | .01 | .97 | 16 | -.13 | .40 | 46 | .19 | .20 | 47 |
| Bulb Volume | Negative | -.21 | .44 | 16 | -.16 | .28 | 46 | -.01 | .97 | 47 |
| Bulb Volume | Disorganized | -.15 | .58 | 16 | -.12 | .44 | 46 | -.12 | .41 | 47 |
| Bulb Volume | General | -.17 | .52 | 16 | -.20 | .17 | 46 | .10 | .49 | 47 |
| Sulcus Depth | Positive | .49 | .05 | 16 | .07 | .63 | 46 | .32 | **.03** | 47 |
| Sulcus Depth | Negative | -.27 | .31 | 16 | -.28 | .06 | 46 | .34 | **.02** | 47 |
| Sulcus Depth | Disorganized | .11 | .69 | 16 | -.18 | .22 | 46 | .24 | .10 | 47 |
| Sulcus Depth | General | -.28 | .30 | 16 | .02 | .87 | 46 | .28 | .06 | 47 |
| Sulcus Length | Positive | .68 | **< .01** | 16 | .09 | .57 | 46 | .16 | .29 | 47 |
| Sulcus Length | Negative | .02 | .93 | 16 | -.31 | **.03** | 46 | .27 | .07 | 47 |
| Sulcus Length | Disorganized | .40 | .13 | 16 | -.05 | .72 | 46 | .21 | .15 | 47 |
| Sulcus Length | General | .08 | .78 | 16 | .02 | .90 | 46 | .09 | .56 | 47 |

*Note. ρ = Spearman rank correlation with age partialled out (residualization method); p = uncorrected p-value; n = valid sample size. Bold p-values indicate p < .05. All analyses are exploratory; findings should be considered hypothesis-generating pending replication. SOPS = Scale of Prodromal Symptoms from the Structured Interview for Prodromal Syndromes subscale totals. Positive direction for odor measures indicates better performance; positive direction for SNOT indicates greater sinonasal symptom burden.*

**Supplemental Table 5. FreeSurfer Euler Number (Scan Quality) by Diagnostic Group**

| **Group** | **n** | **M** | **SD** | **Median** | **Min** | **Max** |
| --- | --- | --- | --- | --- | --- | --- |
| 22q11DS | 24 | -20.3 | 16.4 | -16.0 | -83 | -6 |
| Psychosis Spectrum | 50 | -14.3 | 5.3 | -13.5 | -24 | -4 |
| Typically Developing | 48 | -15.9 | 11.2 | -15.0 | -76 | -2 |

*Note. Mean Euler number averaged across hemispheres, computed from FreeSurfer surface reconstructions of the T1-weighted image (Euler = 2 - 2 x number of topological defect holes; more negative values indicate lower scan quality; Rosen et al., 2018). Sample restricted to participants with usable MRI outcomes (n = 122). Groups did not differ: Kruskal-Wallis H(2) = 1.77, p = .41; all pairwise Mann-Whitney p >= .19, |r| <= .19.*

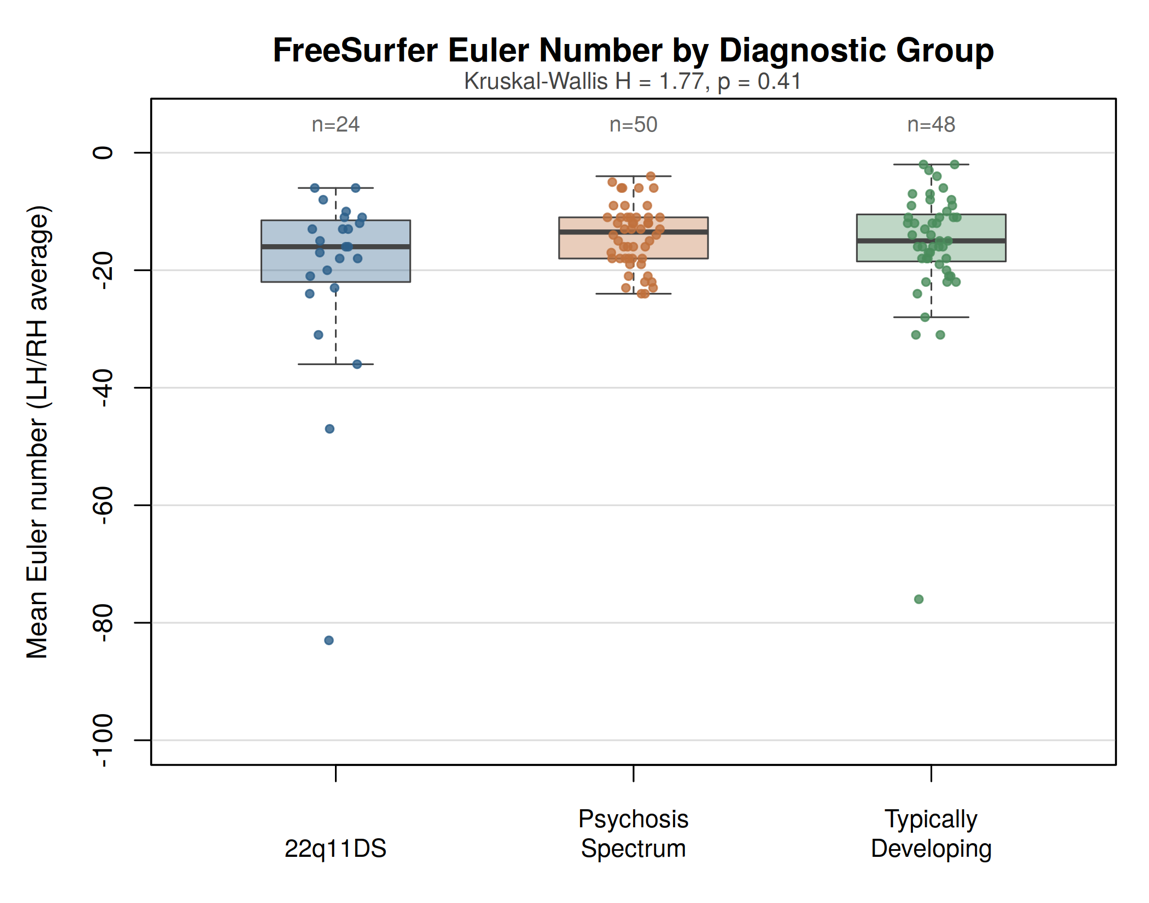

***Supplemental Figure 3. FreeSurfer Euler number by diagnostic group.*** *Distribution of the mean Euler number (averaged across hemispheres) from FreeSurfer reconstruction of the T1-weighted image for participants with usable MRI outcomes. Boxes show median and interquartile range; points are individual participants. More negative values indicate lower surface reconstruction quality. Groups did not differ (Kruskal-Wallis H = 1.77, p = .41).*

**Supplemental Table 6. Covariate-Adjusted Group Contrasts for Primary Olfactory and Sinonasal Outcomes**

| **Measure** | **Contrast** | **Model 1: age + sex** |  |  |  | **Model 2: + race + nonverbal IQ** |  |  |  |
| --- | --- | --- | --- | --- | --- | --- | --- | --- | --- |
|  |  | **n** | **b (SE)** | **p** | **p (Bonf)** | **n** | **b (SE)** | **p** | **p (Bonf)** |
| SNOT | 22q11DS vs TD | 178 | 7.67 (2.43) | 0.002 | 0.006 | 177 | 10.00 (3.27) | 0.003 | 0.008 |
| SNOT | 22q11DS vs PS | 178 | -2.27 (3.20) | 0.480 | 1.000 | 177 | 0.10 (3.53) | 0.977 | 1.000 |
| SNOT | PS vs TD | 178 | 9.94 (2.76) | <.001 | 0.001 | 177 | 9.90 (2.78) | <.001 | 0.001 |
| Odor Identification | 22q11DS vs TD | 106 | -2.47 (0.71) | <.001 | 0.002 | 92 | -3.25 (1.02) | 0.002 | 0.006 |
| Odor Identification | 22q11DS vs PS | 106 | -1.32 (0.79) | 0.096 | 0.287 | 92 | -2.06 (1.01) | 0.043 | 0.130 |
| Odor Identification | PS vs TD | 106 | -1.15 (0.53) | 0.033 | 0.099 | 92 | -1.18 (0.52) | 0.025 | 0.075 |
| Odor Discrimination | 22q11DS vs TD | 81 | -2.98 (0.88) | 0.001 | 0.003 | 73 | -2.68 (1.06) | 0.014 | 0.043 |
| Odor Discrimination | 22q11DS vs PS | 81 | -2.41 (0.81) | 0.004 | 0.012 | 73 | -1.93 (1.14) | 0.095 | 0.285 |
| Odor Discrimination | PS vs TD | 81 | -0.57 (0.60) | 0.340 | 1.000 | 73 | -0.75 (0.61) | 0.225 | 0.674 |
| Olfactory Bulb Volume (mm3) | 22q11DS vs TD | 122 | -18.83 (7.82) | 0.018 | 0.053 | 111 | -14.12 (10.30) | 0.174 | 0.521 |
| Olfactory Bulb Volume (mm3) | 22q11DS vs PS | 122 | -9.71 (8.43) | 0.252 | 0.756 | 111 | -4.33 (9.79) | 0.659 | 1.000 |
| Olfactory Bulb Volume (mm3) | PS vs TD | 122 | -9.12 (5.34) | 0.091 | 0.272 | 111 | -9.78 (5.71) | 0.089 | 0.268 |
| Olfactory Sulcus Depth (mm) | 22q11DS vs TD | 122 | -0.24 (0.45) | 0.594 | 1.000 | 111 | 0.02 (0.68) | 0.972 | 1.000 |
| Olfactory Sulcus Depth (mm) | 22q11DS vs PS | 122 | 0.04 (0.48) | 0.932 | 1.000 | 111 | 0.30 (0.68) | 0.658 | 1.000 |
| Olfactory Sulcus Depth (mm) | PS vs TD | 122 | -0.28 (0.30) | 0.353 | 1.000 | 111 | -0.28 (0.31) | 0.377 | 1.000 |
| Olfactory Sulcus Length (mm) | 22q11DS vs TD | 122 | -0.72 (0.60) | 0.237 | 0.711 | 111 | -0.49 (0.94) | 0.601 | 1.000 |
| Olfactory Sulcus Length (mm) | 22q11DS vs PS | 122 | -0.68 (0.62) | 0.269 | 0.808 | 111 | -0.51 (0.93) | 0.586 | 1.000 |
| Olfactory Sulcus Length (mm) | PS vs TD | 122 | -0.03 (0.45) | 0.941 | 1.000 | 111 | 0.02 (0.47) | 0.972 | 1.000 |

*Note. Ordinary least squares regression of each outcome on group, age (centered, linear and quadratic), and sex (Model 1), additionally adjusting for race (White vs. non-White) and TONI-4 nonverbal IQ (Model 2). b = adjusted group difference (positive values indicate higher scores in the first-listed group); SE and p from heteroscedasticity-consistent (HC3) robust covariance; p (Bonf) = Bonferroni-corrected across three pairwise contrasts per measure. Rank-based ANCOVA on rank-transformed outcomes yielded the same pattern of results. Group x age interactions were non-significant for every outcome (all p >= .13). Adjusting MRI models additionally for the Euler number left all contrasts essentially unchanged, and Euler was not associated with any MRI outcome (all robust p >= .42). Adjustment for nonverbal IQ, a core feature of 22q11DS, can produce biased group estimates (Dennis et al., 2009) and Model 2 is reported as a sensitivity analysis only. Adding current tobacco use to Model 2 left all group contrasts essentially unchanged, and tobacco use was not associated with any outcome (Supplemental Table 11).*

**Supplemental Table 7. Group Comparisons Restricted to the Overlapping Age Range (14-34 Years)**

| **Measure** | **KW H (p)** | **Contrast** | **n** | **r** | **p** | **p (Bonf)** |
| --- | --- | --- | --- | --- | --- | --- |
| SNOT | 21.32 (<.001) | 22q11DS vs TD | (60, 51) | -0.36 | 0.001 | 0.003 |
|  |  | 22q11DS vs PS | (60, 54) | 0.15 | 0.173 | 0.519 |
|  |  | PS vs TD | (54, 51) | -0.51 | <.001 | <.001 |
| Odor Identification | 16.34 (<.001) | 22q11DS vs TD | (20, 42) | 0.61 | <.001 | <.001 |
|  |  | 22q11DS vs PS | (20, 33) | 0.42 | 0.011 | 0.034 |
|  |  | PS vs TD | (33, 42) | 0.25 | 0.067 | 0.200 |
| Odor Discrimination | 13.36 (0.001) | 22q11DS vs TD | (14, 36) | 0.64 | <.001 | 0.001 |
|  |  | 22q11DS vs PS | (14, 28) | 0.57 | 0.003 | 0.008 |
|  |  | PS vs TD | (28, 36) | 0.11 | 0.448 | 1.000 |
| Olfactory Bulb Volume (mm3) | 6.76 (0.034) | 22q11DS vs TD | (19, 48) | 0.37 | 0.018 | 0.054 |
|  |  | 22q11DS vs PS | (19, 50) | 0.19 | 0.222 | 0.665 |
|  |  | PS vs TD | (50, 48) | 0.21 | 0.077 | 0.232 |
| Olfactory Sulcus Depth (mm) | 1.25 (0.536) | 22q11DS vs TD | (19, 48) | 0.12 | 0.453 | 1.000 |
|  |  | 22q11DS vs PS | (19, 50) | 0.01 | 0.973 | 1.000 |
|  |  | PS vs TD | (50, 48) | 0.12 | 0.298 | 0.893 |
| Olfactory Sulcus Length (mm) | 1.49 (0.476) | 22q11DS vs TD | (19, 48) | 0.17 | 0.287 | 0.862 |
|  |  | 22q11DS vs PS | (19, 50) | 0.18 | 0.245 | 0.736 |
|  |  | PS vs TD | (50, 48) | -0.01 | 0.901 | 1.000 |

*Note. Nonparametric analyses (Kruskal-Wallis; pairwise Mann-Whitney U with rank-biserial r) repeated after restricting all groups to ages 14-34 years, the range common to all three groups (22q11DS n = 74 of 101 retained). p (Bonf) = Bonferroni-corrected across three contrasts per measure.*

**Supplemental Table 8. Lifetime Consensus Diagnoses by Diagnostic Group**

| **Lifetime diagnosis category** | **22q11DS (n=101)** | **PS (n=56)** | **TD (n=52)** |
| --- | --- | --- | --- |
| *Primary classification* | | | |
| Threshold psychotic disorder | 11 | 12 | - |
| Psychosis-risk (prodromal) | 56 | 44 | - |
| Typically Developing | - | - | 52 |
| *Secondary lifetime diagnoses (remaining participants)* | | | |
| Bipolar disorder | 0 | 2 | 0 |
| Depressive disorder | 1 | 8 | 4 |
| Mood disorder NOS | 1 | 0 | 0 |
| Anxiety/OCD/PTSD | 14 | 0 | 2 |
| ADHD/disruptive | 3 | 0 | 0 |
| Substance use | 0 | 0 | 3 |

*Note. Diagnoses were assigned by scanning all DSM Axis I codes across each participant's consensus diagnosis records in the laboratory database (median 3 records per participant, 2008-2026). Group membership in the present study was determined by genotype (22q11DS) or SOPS symptom criteria (PS), not by these categorical diagnoses. For 22q11DS, entries in all rows are lifetime primary consensus diagnoses assigned by taking the highest-priority category in a fixed hierarchy (psychotic disorder through no diagnosis), since group membership is defined by genotype irrespective of psychiatric status. For PS and TD, the upper rows show the primary classification and the lower rows show the secondary lifetime diagnoses of participants not represented in the upper rows, so each participant is counted once. All 56 PS participants met SOPS psychosis-spectrum criteria at the time of the study. The 10 PS participants shown under bipolar disorder (n = 2) or depressive disorder (n = 8) carried prodromal codes in their consensus records but did not meet the positive prodromal specifier criteria that define the psychosis-risk row. Their mood diagnoses were documented before study enrollment and several were in remission at the time of the study. Threshold psychotic disorder includes schizophrenia, schizoaffective disorder, delusional disorder, psychotic disorder NOS, and mood disorder with psychotic features (the latter n = 2, both in PS). Psychosis-risk includes positive prodromal or clinical-risk diagnoses (attenuated positive, negative/disorganized, or undifferentiated prodromal syndromes, including remitted). PS classifications were verified and completed against the full clinical database. No TD participant had an active Axis I diagnosis at the time of the study. All secondary lifetime codes in TD derive from database records outside the study period, and all four TD depressive diagnoses were coded in full remission.*

**Supplemental Table 9. Documented Psychotropic and Antipsychotic Medication by Diagnostic Group**

| **Exposure** | **22q11DS** | **PS** | **TD** | **Test** |
| --- | --- | --- | --- | --- |
| Any psychotropic, ever | 58/92 (63.0%) | 27/56 (48.2%) | 5/51 (9.8%) | χ2 = 37.82, p < .001 |
| Antipsychotic, ever | 13/92 (14.1%) | 9/56 (16.1%) | 0/51 (0.0%) | χ 2=8.66, p=0.013 |
| Any psychotropic, current | 57/92 (62.0%) | 24/56 (42.9%) | 4/51 (7.8%) | χ 2 = 39.27, p < .001 |
| Antipsychotic, current | 13/92 (14.1%) | 8/56 (14.3%) | 0/51 (0.0%) | χ 2=8.09, p=0.017 |

*Note. Denominators are participants with any medication record in the laboratory database (199 of 209; 22q11DS 92, PS 56, TD 51). Medications classified by name; psychotropic = antipsychotic, antidepressant, ADHD medication (stimulant or alpha-agonist), anxiolytic/benzodiazepine, or mood stabilizer (lithium, valproate, lamotrigine, oxcarbazepine, carbamazepine). Melatonin and anticonvulsants prescribed predominantly for epilepsy, sleep, or migraine (gabapentin, topiramate, levetiracetam, zonisamide, phenytoin) were not counted. "Current" = record flagged current at any collection; medication collection dates could not be tied precisely to the olfactory session, so exposure is descriptive. Within both clinical groups, antipsychotic-exposed participants reported higher SNOT scores than unexposed participants (22q11DS: M = 27.3 vs. 13.6, p = .008, r = .58; PS: M = 28.4 vs. 16.2, p = .009, r = .58); no behavioral or MRI olfactory measure differed by antipsychotic exposure (all p >= .13).*

**Supplemental Table 10. Olfactory Measures by Psychosis-Spectrum Status Within 22q11DS, and Group Comparisons Excluding 22q11DS+ Participants**

| **Measure** | **22q11DS+ n, M** | **22q11DS− n, M** | **p** | **r** | **22q11DS− vs TD r (corrected p)** | **22q11DS− vs PS r (corrected p)** |
| --- | --- | --- | --- | --- | --- | --- |
| SNOT | 49, 17.2 | 19, 9.8 | .050 | .31 | .18 (.746) | .33 (.102) |
| Odor Identification | 22, 9.4 | 8, 8.8 | .380 | .22 | .76 (.002) | .58 (.035) |
| Odor Discrimination | 14, -- | 2, -- | n/a | n/a | n/a (n = 2) | n/a (n = 2) |
| Bulb Volume (mm3) | 18, 86.1 | 5, 89.1 | .970 | .02 | .48 (.257) | .26 (1.00) |
| Sulcus Depth (mm) | 18, 9.0 | 5, 10.3 | .168 | .42 | .55 (.139) | .59 (.094) |
| Sulcus Length (mm) | 18, 9.2 | 5, 10.8 | .391 | .27 | .24 (1.00) | .21 (1.00) |

*Note. 22q11DS+ = lifetime consensus diagnosis of a psychotic disorder or a positive psychosis-risk (prodromal) syndrome (67 of 91 22q11DS participants with diagnostic data, 73.6%, including 11 with a threshold psychotic disorder); 22q11DS− = neither. This diagnosis-based definition is related to, but not identical to, the SOPS symptom criteria used to define the PS group. Left columns: Mann-Whitney comparisons of 22q11DS+ vs 22q11DS− (rank-biserial r, uncorrected p). Right columns: pairwise contrasts from the three-group analyses repeated after excluding all 22q11DS+ participants (and those without diagnostic data), Bonferroni-corrected. The odor identification deficit remained large and significant in 22q11DS− relative to both TD and PS. Odor discrimination could not be evaluated (only two 22q11DS− participants had discrimination data). MRI contrasts rest on five 22q11DS− participants and are reported for completeness only.*

***Supplemental Table 11. Self-Reported Tobacco Use by Diagnostic Group***

| ***Measure*** | ***22q11DS*** | ***PS*** | ***TD*** | ***Test*** |
| --- | --- | --- | --- | --- |
| *Ever tried tobacco, n (%)* | *8/62 (12.9%)* | *20/51 (39.2%)* | *12/49 (24.5%)* | *Fisher's exact p = .005* |
| *Tobacco use in past 12 months, n (%)* | *1/62 (1.6%)* | *10/51 (19.6%)* | *4/49 (8.2%)* | *Fisher's exact p = .004* |

*Note. Self-reported tobacco use (cigarettes, cigars/pipe, or chewing tobacco) from the study self-report battery, available for 162 of 209 participants (22q11DS 62/101, PS 51/56, TD 49/52). The most severe report was used for the one participant with discordant repeat administrations. Use was infrequent: among the 15 past-12-month users, only one reported daily smoking and most reported use less than once per month. Tobacco use was lowest in 22q11DS, the group with the largest olfactory deficits, so nicotine exposure cannot account for the 22q11DS findings. Repeating the primary group comparisons after excluding all past-12-month users left every significant effect intact (odor identification 22q11DS vs. TD corrected p < .001, vs. PS corrected p = .039; odor discrimination 22q11DS vs. TD corrected p = .002, vs. PS corrected p = .009), and the 22q11DS vs. TD bulb volume reduction was significant in this subsample (corrected p = .012). Adding current tobacco use to the fully adjusted regression models (Supplemental Table 6, Model 2) left all group contrasts essentially unchanged, and tobacco use was not significantly associated with any olfactory outcome (all p >= .06). Fisher's exact test used because expected cell counts were below 5.*
